# Identifying latent genetic interactions in genome-wide association studies using multiple traits

**DOI:** 10.1101/2023.09.11.557155

**Authors:** Andrew J. Bass, Shijia Bian, Aliza P. Wingo, Thomas S. Wingo, David J. Cutler, Michael P. Epstein

## Abstract

Genome-wide association studies of complex traits frequently find that SNP-based estimates of heritability are considerably smaller than estimates from classic family-based studies. This ‘missing’ heritability may be partly explained by genetic variants interacting with other genes or environments that are difficult to specify, observe, and detect. To circumvent these challenges, we propose a new method to detect genetic interactions that leverages pleiotropy from multiple related traits without requiring the interacting variable to be specified or observed. Our approach, Latent Interaction Testing (LIT), uses the observation that correlated traits with shared latent genetic interactions have trait variance and covariance patterns that differ by genotype. LIT examines the relationship between trait variance/covariance patterns and genotype using a flexible kernel-based framework that is computationally scalable for biobank-sized datasets with a large number of traits. We first use simulated data to demonstrate that LIT substantially increases power to detect latent genetic interactions compared to a trait-by-trait univariate method. We then apply LIT to four obesity-related traits in the UK Biobank and detect genetic variants with interactive effects near known obesity-related genes. Overall, we show that LIT, implemented in the R package `lit`, uses shared information across traits to improve detection of latent genetic interactions compared to standard approaches.

## 1 Introduction

There are many large genome-wide association studies (GWAS) available to help facilitate gene discovery and improve our molecular understanding of complex traits and diseases. In particular, recent biobank-sized GWAS collect massive sample sizes and a broad range of phenotypic data to enhance complex trait mapping and augment knowledge of gene functionality. There are two important patterns that have emerged among the multitude of GWAS analyses that have been performed to date. First, the genetic variation of complex traits often involves many thousands of loci [1]. Second, for many traits studied, family-based estimates of heritability (i.e., the proportion of trait variance explained by genetic factors) tend to be substantially greater than corresponding heritability estimates from GWAS single nucleotide polymorphisms (SNP) data [2]. For example, the heritability estimates for body mass index (BMI) from GWAS-based studies are 22−30% [3–6] compared to 40−70% [7,8] in family-based studies. While this ‘missing’ heritability may be due to small sample sizes, structural variants, and/or rare variants [6, 9–11], these sources may not fully explain the difference in some traits [12]. Another possible explanation is that family-based estimates of broad-sense heritability are capturing within-family sharing of genetic variants with interactive effects (e.g., gene-by-gene and gene-by-environment interactions) which are omitted from GWAS estimates derived from nearly unrelated individuals who only generally share the additive effects of alleles (i.e., narrow-sense heritability) [12–14]. Given the evidence of such interactions [15–18], discovering genetic variants with interactive effects may explain missing heritability and broaden our understanding of the genetic architecture of complex traits.

There are many statistical challenges to discovering genetic variants with interactive effects in GWAS [19]. In particular, studies are typically underpowered to detect interactions due to small effect sizes, a large multiple testing burden, and unknown interactive variables (e.g., other variants or environmental factors). Consequently, it can be difficult to design a study to identify and accurately observe interacting variables. Furthermore, even when interacting variables are known, the mismeasurement of such variables can lead to power loss [20]. One strategy to circumvent some of these issues is to use variance-based testing procedures which do not require the interactive variable(s) to be observed [17, 21–24]. Intuitively, such procedures model and detect any unequal residual trait variation among genotype categories at a specific SNP (i.e., heteroskedasticity), which provides evidence of other latent (unobserved) factors interacting with the SNP to influence the trait. Previous work has found that variance-based testing procedures can help identify latent genetic interactions on complex traits, including inflammatory markers [22] and obesity-related traits [17, 23, 24].

When there are multiple related traits measured in a study, researchers often apply variance-based procedures on a trait-by-trait (or univariate) basis to detect latent genetic interactions. However, a univariate strategy ignores any biological pleiotropy among traits despite theoretical [25] and empirical [26, 27] support for this phenomenon. Furthermore, in the presence of pleiotropy, many studies have demonstrated that joint statistical modeling of related traits (i.e., a multivariate procedure) outperforms univariate procedures for gene mapping [28, 29]. This observation, coupled with the potential existence of pleiotropic interactive effects, suggests that analyzing multiple traits simultaneously in a statistical procedure will increase power to detect latent genetic interactions. To this end, we propose a novel statistical framework, called Latent Interaction Testing (LIT), that leverages multiple related traits to increase the power for detecting latent genetic interactions. LIT is motivated by the observation that latent genetic interactions induce not only a differential variance pattern (i.e., heteroskedasticity) as previously reported, but also a differential covariance pattern between traits. We can harness the differential covariance patterns to increase the power to detect latent genetic interactions compared to variance-based strategies. Similar to variance-based strategies, LIT does not require the interactive partner(s) to be observed or specified.

The manuscript is outlined as follows. We first introduce the LIT framework for detecting latent genetic interactions and then evaluate the performance using simulated biobank-sized datasets. We also compare LIT to univariate testing procedures and observe that LIT provides significant power gains to detect interactive effects in GWAS. Finally, we demonstrate LIT using four obesity-related traits in the UK Biobank with over 6 million single nucleotide polymorphisms from 330,868 genotyped individuals. Our analysis identifies multiple loci demonstrating significant interactive effects near known obesity-related genes. We then conclude with a discussion of our results.

## 2 Results

### 2.1 Overview

We provide a brief overview of the LIT framework and leave more technical details to the Methods section. As illustrated in Figure S1, a SNP with a latent interaction induces a genotype effect on the trait variances and on the covariance between traits. We can assess this interaction effect by relating an individual’s genotype to individual-specific trait variances and covariances (Figure S2). We estimate individual-specific trait variances using squared residuals (SQ) for each trait after adjusting for additive (and possibly dominance) effects and likewise estimate individual-specific covariances by multiplying the residuals of different pairs of traits together to form cross products (CP; see ref. [30]). Using a kernel-based distance covariance (KDC) statistic (Figure 1) [31–33], we then assess evidence of a latent genetic interaction by testing whether the elements of a matrix comprised of pairwise similarity of SQ/CP terms in the sample is independent of the elements of a second matrix comprised of pairwise genotype similarity. To measure the similarity between variables, we apply a user-defined kernel function such as a linear kernel (analogous to scaled covariance) or a projection kernel [34, 35]. We show later that the optimal kernel choice depends on the complexity of the interaction signal. Researchers have previously applied variations of the KDC statistic, which yields a *p*-value testing the global null of no association between the elements of two matrices, in genetic analyses for studies of both common [34, 36, 37] and rare [35, 38, 39] variation.

**Figure 1:**
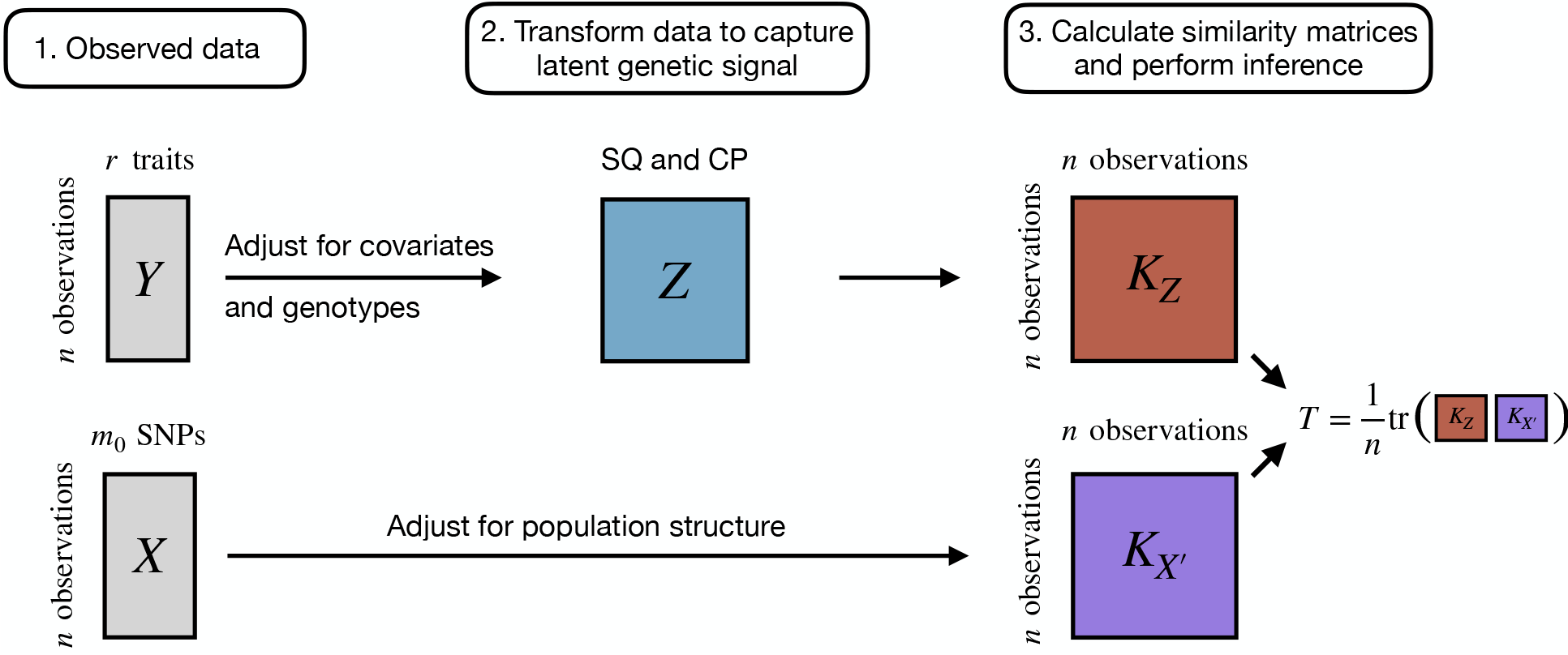
Overview of the Latent Interaction Testing (LIT) framework. Given a set of *r* traits, ***Y***, and *m*_0_ SNPs, ***X***, the goal is to detect a latent genetic interaction involving the SNPs. The trait squared residuals (SQ) and cross products (CP), ***Z***, are calculated while adjusting for linear effects from the genotypes and any other covariates. The traits and genotypes are also adjusted for population structure. A similarity matrix for the genotypes, ***K***_*X′*_, and the SQ and CP, ***K***_*Z*_, are calculated to construct a test statistic, *T*, which measures the overlap between the two matrices. Large values of *T* are evidence of a latent genetic interaction.

The traditional KDC statistic utilizes the corresponding eigenvectors (directions of maximal variation) and eigenvalues (weights emphasizing eigenvectors) derived from the SQ/CP similarity matrix for inference. In the process, the traditional KDC statistic emphasizes signals explaining the most variation in this matrix. While we show this emphasis is suitable under certain pleiotropy settings, there are other settings where the interaction signal is not captured by the top eigenvectors of the similarity matrix and so the test may not be optimal [29, 40]. Therefore, we also consider weighting eigenvectors equally in our test statistic to increase power to detect interaction signals captured by the lower eigenvectors of the similarity matrix. We refer to the implementation that weights eigenvectors by corresponding eigenvalues (i.e., the traditional KDC framework) as weighted LIT (wLIT) and refer to the implementation that weights eigenvectors equally as unweighted LIT (uLIT). Since the pleiotropic genetic architecture of a trait is unknown *a priori*, we maximize the performance of LIT by aggregating the *p*-values from wLIT and uLIT using the Cauchy combination test (CCT) [41], which has proven valuable in a variety of genetic settings [42]. We refer to the CCT of the wLIT and uLIT *p*-values as aggregate LIT (aLIT). For simplicity, we primarily focus on implementing wLIT, uLIT, and aLIT on a SNP-by-SNP basis to test for interactive effects but discuss extensions to handling multiple SNPs simultaneously within the Methods section (see Section 4.2.2). Finally, to improve computational efficiency for biobank-sized datasets, we apply a linear kernel in wLIT and so uLIT is equivalent to using a projection kernel.

### 2.2 Power and type I error rate control

We simulated *r* = 5, 10 related traits for 300,000 observations (reflecting sample sizes for biobank datasets) under the polygenic trait model with additive genetic, environmental, and GxE interaction components. The baseline correlation between traits was either 0.25, 0.50, or 0.75 which represents different correlation strengths from shared genetic and environmental effects. We then simulated a genetic risk factor, an environmental factor, and a GxE interaction that explains 0.2%, a randomly drawn value from 0.5% to 2.0%, and a randomly drawn value from 0.1% to 0.15% of the trait variation, respectively. To assess the performance of LIT under different sparsity settings, we varied the proportion of traits with a shared GxE interaction as 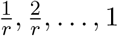. We considered three types of pleiotropy in our study, namely, the GxE interaction effect size is positive across traits (positive pleiotropy), a mixture of positive and negative across traits (positive and negative pleiotropy), and a variation of these two settings where the direction is opposite of the interacting environment.

We found that the LIT implementations provide type I error rate control at significance level 10^−3^, including when the trait distribution is skewed or heavy trailed (Figure S3,S4). We then compared the power across various configurations of number of traits, baseline correlation, proportion of traits with shared interaction effects, and direction of the interaction effect (Figure 2A-B). In comparing wLIT with uLIT, neither method is optimal across all settings as expected. As mentioned in Section 2.1, wLIT emphasizes the high-variance (i.e., large eigenvalues) eigenvectors of the SQ/CP kernel matrix while uLIT weights them equally. Under a simulation model where the signal would reside on the top eigenvector, we expect wLIT to outperform uLIT. Conversely, we expect uLIT to outperform wLIT when the signal resides on the lower-variance eigenvectors of the SQ/CP kernel matrix.

**Figure 2:**
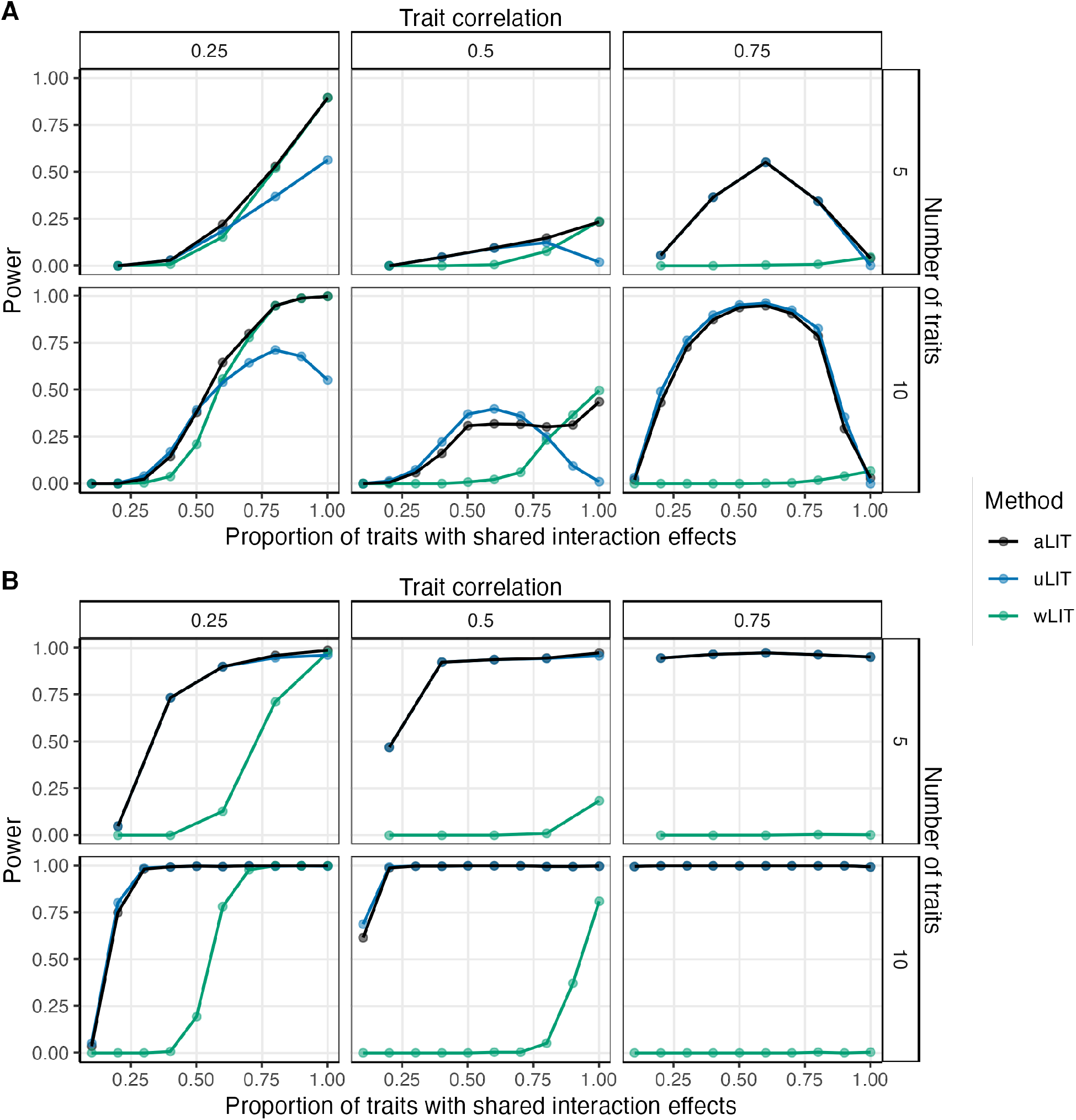
Power comparisons of aggregate LIT (aLIT; black), unweighted LIT (uLIT; blue), and weighted LIT (wLIT; green) under (**A**) positive pleiotropy and (**B**) a mixture of positive and negative pleiotropy. The simulation study varied the correlation between traits (columns), the number of traits (rows), and the proportion of traits with an interaction term (x-axis) at a sample size of 300,000. The points represent the average across 500 simulations with a significance threshold of 5 *×* 10^−8^.

To illustrate how the interaction signal can reside on different eigenvectors of the SQ/CP kernel matrix, we performed an association test between the eigenvectors and genotype under the positive pleiotropy setting with 10 traits (Figure S5). We find that the power to detect the latent interaction signal at each eigenvector depends on the proportion of traits with shared interaction effects (sparsity level), baseline trait correlation, and the proportion of variation explained by the genotype (denoted as *R*^2^). More specifically, for small baseline correlations, the high-variance eigenvector generally captures the signal for most sparsity settings (which explains why wLIT outperforms uLIT in these situations). As the baseline correlation increases, the power of the high-variance eigenvector can decrease rapidly (even if the proportion of traits with shared interaction effects is high) due to the reduction in *R*^2^ (see top right panels of Figure S5). On the other hand, an increase in baseline correlation coupled with a decrease in the proportion of traits with shared interaction effects (i.e., increase in sparsity), results in the interaction signal being separated out from the high-variance eigenvector and becoming detectable in the low-variance eigenvectors.

Given the above insights, we can delineate the performance between uLIT and wLIT by assessing which eigenvectors capture the interaction signal. In the positive pleiotropy setting with 10 traits and a baseline correlation of 0.5 (bottom center panel of Figure 2A), as the proportion of traits with shared interaction effects increases from 0 to 0.6, the power of uLIT increases whereas the power of wLIT is constantly negligible. When the proportion of traits with shared interaction effects increases from 0.6 to 1, the power of uLIT decreases whereas the power of wLIT increases and overtakes uLIT when this proportion exceeds 0.8. These power trends are due to the low-variance eigenvectors capturing the interaction signal when the proportion of traits with shared interaction effects is small (which favors uLIT) to the high-variance eigenvectors when this is high (which favors wLIT; Figure S5). Furthermore, when the baseline correlation increases from 0.5 to 0.75 (bottom right panel of Figure 2A), uLIT follows a similar power curve while the power of wLIT now remains negligible across all sparsity settings. In this case, the *R*^2^ is low in the high-variance eigenvectors when the baseline correlation is high (Figure S5) and so wLIT has little power in these situations. In general, we find that wLIT tends to outperform uLIT when the baseline correlation is modest (i.e., 0.25) and the proportion of traits with shared interaction effects is high, otherwise uLIT is the optimal method.

In the setting where there is a mixture of positive and negative pleiotropy, uLIT outperforms wLIT across all settings (Figure 2B). Intuitively, in our simulations, the high-variance eigenvector is the weighted sum of the squared residuals and cross products where the weights have the same sign. When the effect sizes are in different directions (positive and negative pleiotropy), the high-variance eigenvector may dampen the interaction signal, and thus it will also be captured by the low-variance eigenvectors. Since uLIT weights the eigenvectors equally, we observe a large increase in power compared to wLIT. We also considered a variation of the above two pleiotropy scenarios where the effect size for the GxE interaction is opposite of the interacting environment. While we find similar results, the overall power is reduced for all methods (Figure S6).

In summary, even though the top eigenvectors explain the largest amount of variation, it does not imply that they are the ones most correlated to genotype. The interaction signal may be captured by the high-variance eigenvectors or the low-variance eigenvectors depending on the number of traits, baseline correlation, *R*^2^ at each eigenvector, proportion of traits with shared interaction effects, and type of pleiotropy. Since the particular eigenvectors that are most powerful can vary widely and are unknown *a priori*, we applied aLIT to the *p*-values from the above LIT implementations to maximize the number of discoveries. We find that aLIT controls the type I error rate (Figure S3,S4) while making more discoveries than each individual implementation (Figure 2,S6). More specifically, aLIT has similar power to wLIT when the signal is captured by the high-variance eigenvectors and similar power to uLIT when the signal is captured by the low-variance eigenvectors. Therefore, we implement aLIT in subsequent analyses.

### 2.3 aLIT increases power compared to marginal testing procedures

Using the same simulation configuration as in Section 2.2, we considered two competing procedures for identifying latent genetic interactions using multiple traits. The first procedure performs an association test between the squared residuals and a SNP (Marginal (SQ)), while the second procedure additionally includes the cross product terms (Marginal (SQ/CP)). More specifically, Marginal (SQ) tests the squared residuals for all *r* traits and selects the minimum *p*-value from these *r* different tests. Marginal (SQ/CP) adds tests for the 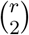 cross products and selects the minimum *p*-value from the 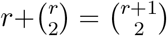 individual tests. Because we are testing the global null hypothesis of no latent genetic interaction, the marginal testing procedures require a Bonferroni correction for the total number of tests, i.e., *α*^*′*^ := *α/K* where *α* is the significance threshold and *K* is chosen to be the number of principal components that explains 95% of the variation. We then threshold the minimum *p*-value by *α*^*′*^ to determine statistical significance. Across all power simulations (Figure 3, S7), we observed that Marginal (SQ/CP) was more powerful than Marginal (SQ), suggesting that the inclusion of cross products improves performance to detect latent interactions. Given these findings, we compare the performance of aLIT to Marginal (SQ/CP) for the remainder of this work.

**Figure 3:**
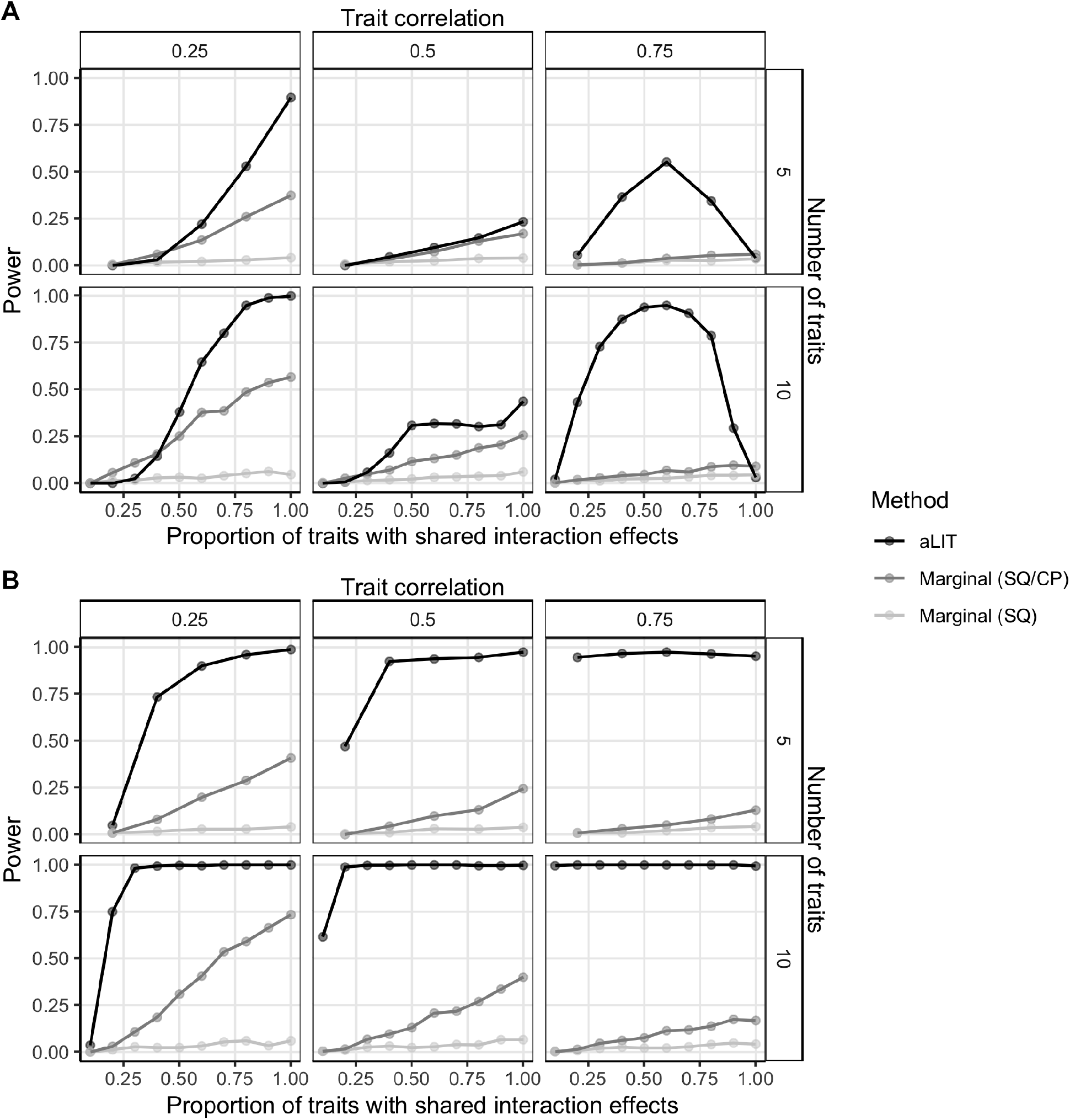
A comparison of aLIT (black) to a marginal testing procedure using the squared residuals (SQ; light grey) and a marginal testing procedure using the squared residuals and cross products (SQ/CP; dark grey) under (**A**) positive pleiotropy and (**B**) a mixture of positive and negative pleiotropy. The simulation study is identical to Figure 2 where the empirical power is calculated as a function of the proportion of traits with an interaction term (x-axis), the number of traits (rows), and trait correlation (columns).

In the positive pleiotropy setting with a low baseline correlation, aLIT increases the power to detect GxE interactions when there are a higher proportion of traits with shared interaction effects compared to Marginal (SQ/CP) (Figure 3A). Furthermore, the difference is more pronounced as the number of traits increases. For example, when the baseline correlation is 0.25, and the proportion of traits with shared interaction effects is 0.8, the empirical power of aLIT is 53% and 94.8% for five and ten traits, respectively. On the other hand, the empirical power of Marginal (SQ/CP) is 26% and 48.6%, respectively. While aLIT provides substantial increases in power when the proportion of traits with shared interaction effects is high, Marginal (SQ/CP) can outperform aLIT when the proportion of traits with shared interaction effects is low in the positive pleiotropy setting (see lower left panel of Figure 3A). Intuitively, when there is little correlation between traits due to shared interactions, selecting the minimum *p*-value across traits slightly outperforms combining information between the traits.

The difference in power between aLIT and Marginal (SQ/CP) is also evident across baseline correlations. Interestingly, the improvement in power of aLIT compared to Marginal (SQ/CP) reduces when the baseline correlation increases from 0.25 to 0.50. This observation agrees with our simulation results from Figure S5 which suggest that the power to detect an interaction signal at any particular eigenvector decreases as the baseline correlation increases from 0.25 to 0.50. Alternatively, when the baseline correlation increases to 0.75, aLIT provides drastic increases in power for most sparsity settings. For example, when there are 10 traits where the proportion of traits with shared interaction effects is 0.5, aLIT’s power increases from 30.8% to 93.8% at a baseline correlation of 0.50 and 0.75 while Marginal (SQ/CP) decreases from 11.6% to 4.6%, respectively. However, in this same example, Marginal (SQ/CP) slightly outperforms aLIT when all traits have an interaction.

Overall, while our results suggest that aLIT outperforms Marginal (SQ/CP) for most baseline correlations and sparsity settings under positive pleiotropy, there are some rare cases where Marginal (SQ/CP) has similar (or improved) performance. Meanwhile, under the simulation setting where there is a mixture of positive and negative pleiotropy (Figure 3B) or the direction of GxE effect sizes are opposite of the interactive environment (Figure S7), the increase in power from aLIT over Marginal (SQ/CP) is substantial across all settings. Note that aLIT outperformed Marginal (SQ) across all power simulations.

### 2.4 aLIT applied to the UK Biobank data

We applied the LIT framework to detect shared latent genetic interactions in four obesity-related traits from the UK Biobank, namely, waist circumference (WC), hip circumference (HC), body mass index (BMI), and body fat percentage (BFP). After preprocessing, there were 329,146 unrelated individuals that have measurements for all traits and 6,186,503 SNPs (Section 4.4). The correlation between traits ranged from 0.75 (BMI and BFP) to 0.87 (BMI and WC). The total computational time of LIT was approximately 3.3 days using 12 cores.

In each implementation of LIT, after filtering for LD and significant SNPs, the genomic inflation factor of wLIT and uLIT was 1.14 (Figure S8a). While the test statistics are inflated, it is difficult to distinguish the factors driving inflation, e.g., unmodeled population structure or biological signal under polygenic inheritance [43]. We found that the genomic inflation factor increases as a function of minor allele frequency which is expected under polygenic inheritance (Figure S9). To be conservative, we adjusted the significance results of each approach by the corresponding genomic inflation factor (Figure S8b). These adjusted *p*-values were then combined in aLIT to detect latent genetic interactions (Figure 4).

**Figure 4:**
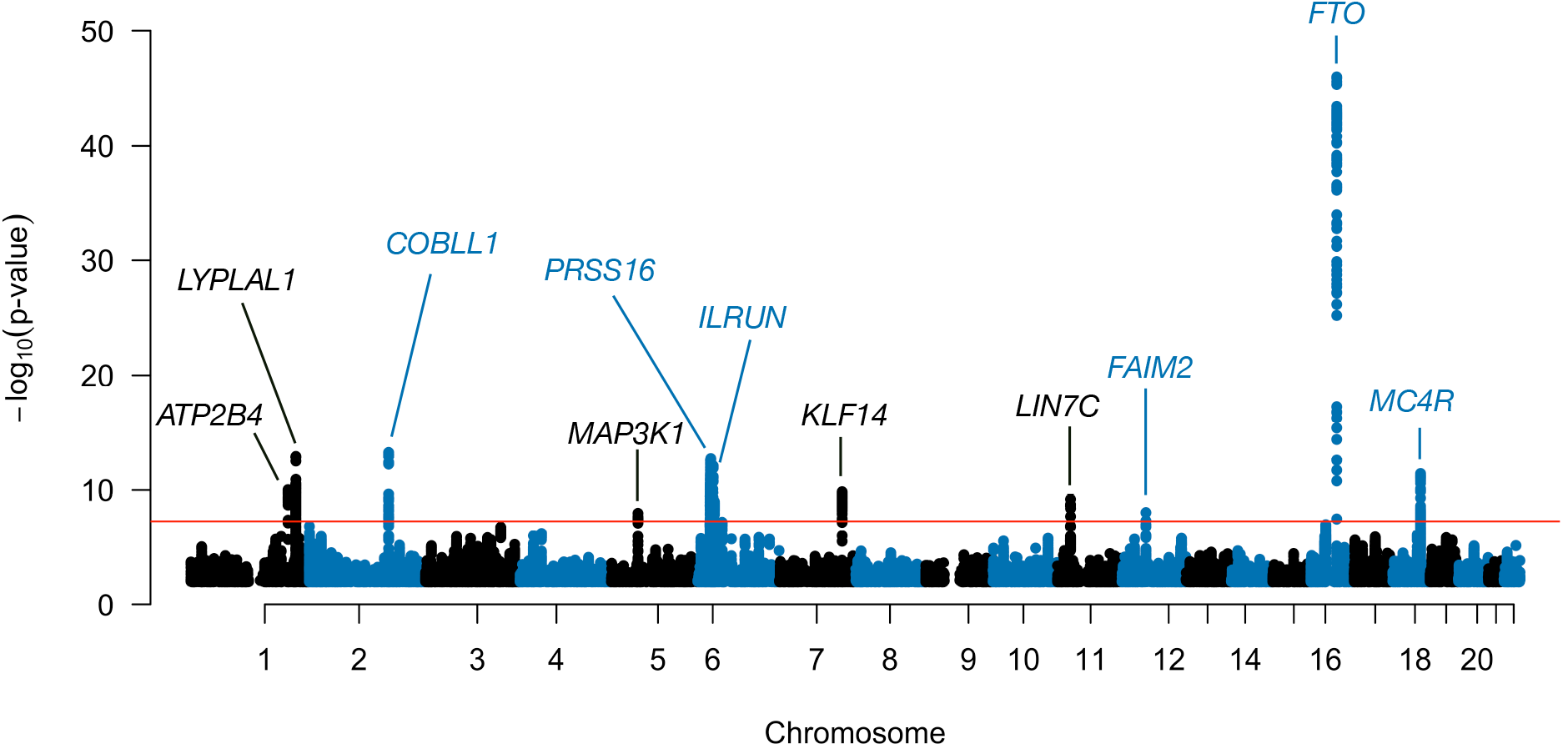
Manhattan plot of aLIT *p*-values using obesity-related traits (waist circumference, hip circumference, body mass index, and body fat percentage) in the UK Biobank. The red line represents the genome-wide significance threshold of 5 *×* 10^−8^. Note that *p*-values below 0.1 are removed from the plot.

Using the aLIT *p*-values, we discovered 2,252 SNPs with significant interactive effects in 11 distinct regions. Table 1 shows the most significantly associated (lead) SNP in each region. As a comparison, we also applied Marginal (SQ/CP) and detected 2,099 SNPs. Of those found by Marginal (SQ/CP), aLIT’s results overlapped with ≈98% of the detected SNPs and had substantially smaller *p*-values at most loci (Figure S10). Although Marginal SQ/CP detects a few regions that are not found by aLIT, the aLIT *p*-values are comparable in magnitude at these regions. On the other hand, there are three distinct regions found by aLIT (rs2821230, rs11030066, and rs157845) where the *p*-values are substantially smaller than the *p*-values from Marginal (SQ/CP) (Table 1). Thus, in agreement with our simulation results, depending on the type of pleiotropy at a particular locus, there are regions where the significance results of Marginal (SQ/CP) and aLIT are comparable and other regions where aLIT is substantially more powerful than Marginal (SQ/CP).

**Table 1:** aLIT and Marginal (SQ/CP) significance results of the lead SNPs from the UK Biobank analysis. There are two other *p*-values reported to help assess statistical significance in aLIT: (i) accounting for significant SNPs in linkage disequilibrium with the lead SNP (labeled ‘LD’) and (ii) removing dominance and/or scaling effects (labeled ‘Dom.’).

| Chr. | Gene | Lead SNP | MAF | $p$-value (aLIT) | $p$-value (SQ/CP) | $p$-value (LD) | $p$-value (Dom.) |
| --- | --- | --- | --- | --- | --- | --- | --- |
| 16 | *FTO* | rs11642015 | 0.402 | $1.08 \times 10^{-46}$ | $2.73 \times 10^{-40}$ | $1.23 \times 10^{-46}$ | $1.28 \times 10^{-46}$ |
| 2 | *COBLL1* | rs5835988 | 0.406 | $5.30 \times 10^{-14}$ | $1.08 \times 10^{-9}$ | $1.29 \times 10^{-13}$ | $5.32 \times 10^{-14}$ |
| 1 | *LYPLAL1* | rs2820444 | 0.299 | $1.17 \times 10^{-13}$ | $2.64 \times 10^{-13}$ | $1.27 \times 10^{-13}$ | $1.19 \times 10^{-13}$ |
| 6 | *PRSS16* | rs13212921 | 0.136 | $1.78 \times 10^{-13}$ | $3.40 \times 10^{-12}$ | $1.90 \times 10^{-12}$ | $4.85 \times 10^{-9}$ |
| 18 | *MC4R* | rs35614134 | 0.234 | $3.95 \times 10^{-12}$ | $9.34 \times 10^{-12}$ | $1.39 \times 10^{-11}$ | $3.97 \times 10^{-12}$ |
| 1 | *ATP2B4* | rs2821230 | 0.474 | $9.45 \times 10^{-11}$ | $1.19 \times 10^{-5}$ | $1.02 \times 10^{-10}$ | $9.38 \times 10^{-11}$ |
| 7 | *KLF14* | rs972284 | 0.389 | $1.43 \times 10^{-10}$ | $2.43 \times 10^{-9}$ | $1.25 \times 10^{-10}$ | $1.43 \times 10^{-10}$ |
| 11 | *LIN7C* | rs11030066 | 0.143 | $6.38 \times 10^{-10}$ | $5.03 \times 10^{-6}$ | $6.97 \times 10^{-10}$ | $6.62 \times 10^{-10}$ |
| 6 | *ILRUN* | rs9469860 | 0.146 | $1.01 \times 10^{-9}$ | $4.47 \times 10^{-9}$ | $7.72 \times 10^{-9}$ | $3.62 \times 10^{-7}$ |
| 12 | *FAIM2* | rs7132908 | 0.384 | $8.85 \times 10^{-9}$ | $4.22 \times 10^{-9}$ | $8.49 \times 10^{-9}$ | $8.96 \times 10^{-9}$ |
| 5 | *MAP3K1* | rs157845 | 0.253 | $1.03 \times 10^{-8}$ | $2.39 \times 10^{-4}$ | $1.30 \times 10^{-8}$ | $1.03 \times 10^{-8}$ |

A concern in this analysis is whether statistical significance is due to a nearby SNP with a large additive effect and/or trait scaling issues. To address the latter, first note that the LIT framework not only detects latent interactions, but also departures from linearity due to dominance or misspecification of the trait scale. Therefore, to be conservative and help distinguish interactive effects, we removed the non-linear genetic signal within a locus by fitting a two degree of freedom genotypic model (Section 4.5). After removing the dominance/scaling effects, the lead SNP rs9469860 on chromosome 6 was above the genome-wide significance threshold (Table 1). In general, while the significance of a few loci were impacted (nearly all on chromosome 6 and a few on chromosome 18; Figure S11), most of the lead SNPs remained significant. Finally, to account for nearby SNPs with large additive effects, we applied the LIT implementations to the lead SNPs while regressing out nearby significant SNPs in LD and found all of the lead SNPs remain significant (Table 1).

After evaluating the significant SNPs found by aLIT, we focused on ten lead SNPs that remained significant after accounting for LD and dominance/scaling issues. Using the Variants to Genes (V2G) measure on Open Targets Platform [44], we assigned the lead SNPs to the highest ranked genes (Table 1). A few of these genes are found in other GWAS of obesity-related traits. In particular, the *FTO* gene is a known obesity-related gene that is associated with type 2 diabetes (see, e.g., [45–47]). While V2G score assigned rs5935988 to *COBLL1* (198,262 bp), it was also close to *GRB14* (23,935 bp) which has been associated with body fat distribution and may be involved in regulating insulin signaling [48–50]. Other genes that were assigned to lead SNPs are involved in regulating satiety and energy homeostasis (*MC4R*; [51]), adiposity (*LYPLAL1*; [52, 53]), and metabolic diseases such as type 2 diabetes (*KLF14*; [54, 55]).

We then searched for evidence of known interacting variables using the significant SNPs identified by aLIT. Previous work has found sex-specific effects of variants in *KLF14, GRB14*, and *LYPLAL1* [50, 55, 56]. Therefore, we tested whether sex was an interactive variable in at least one of the traits using a multivariate regression model and found evidence of a genotype-by-sex interaction in rs972284 (*p* = 2.39 *×* 10^−14^; *KLF14*), rs5835988 (*p* = 1.32 *×* 10^−51^; *COBLL1*/*GRB14*), and rs2820444 (*p* = 7.23 *×* 10^−23^; *LYPLAL1*).

## 3 Discussion

It is challenging to identify, observe, accurately measure, and then detect genetic interactions in a GWAS study. While there are methods to infer interactions that do not require specifying the interactive partner(s) [17, 22–24], these approaches only consider a single trait. To increase statistical power, our proposed kernel-based framework, Latent Interaction Testing (LIT), leverages the shared genetic interaction signal from multiple related traits (i.e., pleiotropy) while maintaining the flexibility of single trait approaches. In our simulation study, we found that the optimal implementation between wLIT and uLIT depends on the genetic architecture. We also found that combining the *p*-values from both approaches in aLIT maximized the number of discoveries while controlling the type I error rate. Furthermore, aLIT increased the power to detect latent genetic interactions compared to marginal testing procedures, and the difference was drastic for certain genetic architectures. We then applied the LIT framework to four obesity-related traits in the UK Biobank and found many loci with potential interactive effects. While we emphasized the linear and projection kernels in our study, aLIT can incorporate multiple kernel choices (e.g., Gaussian) which may increase the power to detect complex interaction signals. However, including additional kernel functions will also increase the computational complexity which may be time prohibitive for biobank-sized datasets.

There are some caveats when interpreting the significance results of LIT or, more generally, any approach that does not require observing the interactive variable(s). Type I error rate control is impacted by loci with large additive effects and trait scaling issues. To address the former issue, we performed inference using all SNPs in LD with the lead SNPs. While it is possible that the true causal SNP is not tagged, it is unlikely in this work since there is dense coverage with the imputed genotypes. We also assessed model misspecification due to an incorrect trait scaling by fitting a genotypic model to flexibly capture non-linear genetic signals. Interestingly, we found that our significance results were primarily impacted at loci located on chromosome 6 (outside the MHC). While this strategy can help identify the extent of genome-wide inflation due to dominance/scaling, it cannot determine whether the latent interactive effects are an artifact of the scale, even though our simulations suggest that detections by LIT are robust to deviations from normality. When presented with traits that follow a non-Normal distribution, an inverse-Normal transformation is typically applied so a trait “appears” as a standard Normal distribution. However, for detecting latent interactions, we recommend against such practice as it does not correct for the mean-variance relationship and can lead to invalid inference [24]. In general, model misspecification from an incorrect scaling is problematic for any population genetic analysis and may require other approaches such as goodness-of-fit testing to help identify an appropriate variance-stabilizing transformation. Finally, similar to other variance-based testing procedures, LIT cannot distinguish whether a discovery is due to a gene-by-gene interaction, gene-by-environment interaction, parent-of-origin effects, or another complex non-additive relationship involving the tested SNP.

There are also several important considerations when applying LIT to genetic data. Importantly, in this work, we assume that individuals are unrelated and traits follow a multivariate Normal distribution. While LIT assumes the data follows a multivariate Normal distribution, our simulation study suggests that it is robust to violations of this assumption. In general, the computational time of LIT increases as the number of traits and sample size increases (Figure S12). Therefore, in order to analyze biobank-sized datasets, LIT uses multiple cores to distribute SNPs (e.g., on the same chromosome) for interaction testing to be computationally more efficient. Because calculating the residual cross products for a large number of traits is computationally intensive (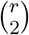 increase in computational time per SNP), LIT provides a user option to only use the squared residuals. However, as demonstrated in simulation and in the UK Biobank dataset, employing this option is nearly certain to lose power. Finally, while a discovery in LIT suggests evidence of a non-additive effect, LIT does not identify the trait, or subset of traits, driving that result. To do so, investigators might consider running Marginal (SQ/CP) at the “lead” SNP to rank/identify individual traits with non-additive effects (squared residuals) and pairs of traits with shared non-additive effects (cross products; although, see Methods).

For many complex traits, there is strong discrepancy between GWAS-based estimates of heritability (which explicitly assume additive effects of genetic variation) and family-based estimates (which may incorporate non-additive effects and higher-order interactions). With recent biobank-sized datasets, we can begin to identify loci with non-additive genetic variation that contribute to this missing heritability while understanding its role in the etiology of complex traits. As biobank-sized datasets become more prevalent, we anticipate that computationally scalable approaches that leverage information across multiple traits, such as LIT, will become increasingly important to discovering non-additive genetic loci.

## 4 Methods

### 4.1 Motivation

Consider the trait *Y*_*jk*_ for *j* = 1, 2, …, *n* unrelated individuals with *k* = 1, 2, …, *r* measurable traits. Suppose *Y*_*jk*_ depends on a biallelic locus with genotype *X*_*j*_ denoting the number of minor alleles for the *j*th individual, an unobserved (or latent) environmental variable *M*_*j*_, and a latent genotype-by-environment (GxE) interaction *X*_*j*_*M*_*j*_. These components contribute to expression additively in the following regression model:

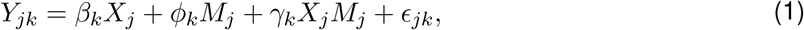

where *β*_*k*_ is the effect size of the minor allele, *ϕ*_*k*_ is the effect size of the environmental variable, *γ*_*k*_ is the effect size of the GxE interaction, and *ϵ*_*jk*_ is an independent and identically distributed random error with mean zero and variance 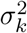. In this simplified setting, our goal is to detect the latent GxE interaction without observing the interacting variable *M*_*j*_.

Under the above model assumptions, the latent GxE interaction will induce differential trait variance and covariance patterns that differ by genotype. Without loss of generality, assume the environmental variable has mean zero with unit variance. In Appendix 5.1.1, we show that the individual-specific trait variance (ITV) of the *k*th trait conditional on genotype is

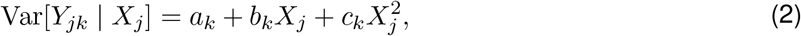

where 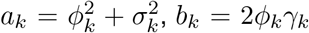, and 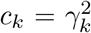. We also show that the individual-specific covariance (ITC) between the *k*th and *k*^*′*^th trait conditional on genotype is

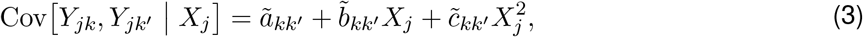

where 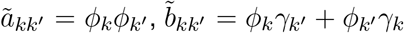, and 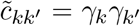. It is evident that a latent GxE interaction in trait *k* (*γ*_*k*_≠ 0) not only induces a variance pattern that depends on genotype (Equation 2), but can also induce a covariance pattern between traits *k* and *k*^*′*^ from either a shared interaction (*γ*_*k′*_≠ 0)or a shared environment involved in the interaction (*ϕ*_*k′*_≠ 0; Equation 3). These results suggest that we can test for loci with latent interactive effects by assessing whether the individual-specific trait variances (ITV) and covariances (ITC) differ by genotype without specifying or directly modeling the interacting variable *M*_*j*_. While we assumed the interacting variable is environmental, the above insights are generalizable to any latent interaction(s) involving the SNP *X*_*j*_ (e.g., a genotype-by-genotype interaction).

### 4.2 Latent Interaction Testing (LIT) framework

Our strategy builds from the above observations and estimates the ITV and ITC to detect latent genetic interactions. To derive estimates of these quantities, we first remove the additive genetic effect from the traits to ensure that any variance and covariance effects are not due to the additive effect. Let us denote the trait residuals as *e*_*jk*_ = *Y*_*jk*_ − *β*_*k*_*X*_*j*_ where we assume the effect size is known for simplicity. We can then express the ITV and ITC as a function of these residuals: the ITV of trait *k* and the ITC between traits *k* and *k*^*′*^ is defined as 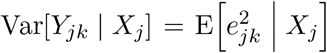 and 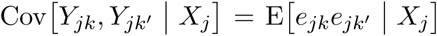, respectively (Appendix 5.1.1). Thus, we can estimate the ITV by squaring the residuals, 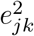, and estimate the ITC between traits *k* and *k*^*′*^ by the pairwise product of the residuals (i.e., the cross products), *e*_*jk*_*e*_*jk′*_. Aggregating the ITV and ITC estimates across all individuals, we denote the cross product (CP) terms in the *n× s* matrix ***Z***^CP^ where the *j*th row vector is 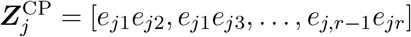, and the squared residual (SQ) terms in the *n × r* matrix ***Z***^SQ^ where 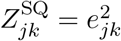.

Our inference goal is to assess whether the SNP, ***X***_*n×*1_ = [*X*_1_, *X*_2_, …, *X*_*n*_]^*T*^, is independent of the squared residuals and cross products,

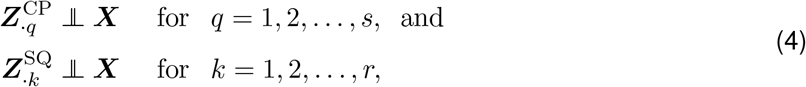

where ‘·’ denotes all the rows (or individuals) and ‘**⫫**’ denotes statistical independence. In the above regression model, this corresponds to testing the global null hypothesis *H*_0_ : *γ*_1_ = *γ*_2_ = … = *γ*_*r*_ = 0 versus the alternative hypothesis *H*_1_ : *γ*_*k*_≠ 0 for at least one of the *k* = 1, 2, …, *r* traits. While a regression model can be directly applied to the squared residuals and cross products to test the global null hypothesis (see Appendix for mathematical details), a univariate model approach does not adequately leverage pleiotropy and requires a multiple testing correction which reduces power.

To address these issues, we develop a new multivariate kernel-based framework, Latent Interaction Testing (LIT), that captures pleiotropy across the ITV and ITC terms to increase power for detecting latent interactions. There are three key steps in the LIT framework (Figure 1):

1. Regress out the additive genetic effects and any other covariates from the traits. Additionally, adjust the traits and genotypes for population structure.
2. Calculate estimates of the ITV and ITC for each individual using the squared residuals and the cross products of the residuals, respectively.
3. Test the global null hypothesis of no latent interaction by comparing the adjusted genotype(s) to the ITV and ITC estimates.

We expand on the above steps in detail below.

**Step 1**: In the first step, LIT standardizes the traits and then regresses out the additive genetic effects, population structure, and any other covariates. This ensures that any differential variance and/or covariance patterns are not due to additive genetic effects or population structure. Suppose there are *l*_1_ measured covariates and *l*_2_ principal components to control for structure. We denote these *l* = *l*_1_ + *l*_2_ variables in the *n × l* matrix ***H***. After regressing out these variables and the additive genetic effects, the *n × r* matrix of residuals is 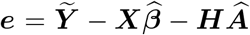, where 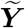 is the standardized trait matrix, 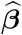 is a 1 *× r* matrix of effect sizes and 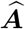 is a *l × r* matrix of coefficients estimated using least squares. We also regress out population structure from the genotypes which we denote by 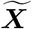.

The above approach only removes the mean effects and does not correct for variance effects from population structure which can impact type I error rate control [57]. A strategy to adjust for the variance effects is to standardize the genotypes with the estimated individual-specific allele frequencies (IAF), i.e., the allele frequencies given the genetic ancestry of an individual. However, it is computationally costly to standardize the genotypes for biobank-sized datasets as it requires estimating the IAFs of all SNPs using a generalized linear model [58, 59]. Therefore, in this work, we remove the mean effects from structure and then adjust the test statistics with the genomic inflation factor to be conservative. Our software includes an implementation to standardize the genotypes using the IAFs for smaller datasets.

**Step 2**: The second step uses the residuals, ***e***, to reveal any latent interactions by constructing estimates of the ITV and ITC. For the *j*th individual’s set of trait residuals, the ITVs are estimated by squaring the trait residuals while the ITCs are estimated by calculating the cross products of the trait residuals. We express the squared residuals as 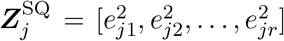, and the 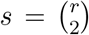 pairwise cross products as 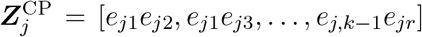. Importantly, when the studentized residuals are used, then 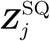 and 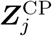 represent an unbiased estimate of the ITVs and ITCs, respectively. We aggregate these terms across all individuals into the *n ×* (*r* + *s*) matrix ***Z*** = [***Z***^SQ^ ***Z***^CP^].

**Step 3**: In the last step, we test for association between the adjusted SNP and the squared residuals and cross products (SQ/CP) using a kernel-based distance covariance framework [31–33]. Specifically, we apply a kernel-based independence test called the Hilbert-Schmidt independence criterion (HSIC), which has been previously used for GWAS data (see, e.g., [35–38]). The HSIC constructs two *n × n* similarity matrices between individuals using the SQ/CP matrix and genotype matrix, then calculates a test statistic that measures any shared signal between these similarity matrices. To estimate the similarity matrix, a kernel function is specified that captures the similitude between the *j*th and *j*^*′*^th individual.

Since our primary application is biobank-sized data, we use a linear kernel so that LIT is computationally efficient. The linear similarity matrix is defined as 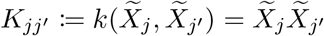 for the genotype matrix and 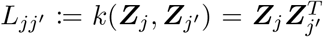 for the SQ/CP matrix. The linear kernel is a scaled version of the covariance matrix and, for this special case, the HSIC is related to the RV coefficient. We note that one can choose other options for a kernel function, such as a polynomial kernel, projection kernel, and a Gaussian radial-basis function that can capture non-linear relationships [34, 35].

Once the similarity matrices ***K*** and ***L*** are constructed, we can express the HSIC test statistic as

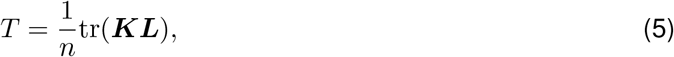

which follows a weighted sum of Chi-squared random variables under the null hypothesis, i.e., 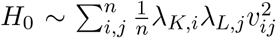, where *λ*_*K,i*_ and *λ*_*L,j*_ are the ordered non-zero eigenvalues of the respective matrices and *v*_*ij*_ ∼ Normal(0, 1). Intuitively, the test statistic measures the ‘overlap’ between two random matrices where large values of *T* imply the two matrices are similar (i.e., a latent genetic interactive effect) while small values of *T* imply no evidence of similarity (i.e., no latent genetic interactive effects). We can approximate the null distribution of *T* using Davies’ method, which is computationally fast and accurate for large *T* [35, 38, 60].

For the linear kernel considered here, we implement a simple strategy to substantially improve the computational speed of LIT. We first calculate the eigenvectors and eigenvalues of the SQ/CP and genotype matrices to construct the test statistic. Since the number of traits, *r*, is much smaller than the sample size, *n*, we can perform a singular value decomposition to estimate the subset of eigenvectors and eigenvalues in a computationally efficient manner [61–63]. This allows us to circumvent direct calculation and storage of large *n × n* similarity matrices. Let 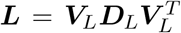 and 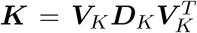 be the singular value decomposition (SVD) of the similarity matrices where the matrix ***D*** is a diagonal matrix of eigenvalues and ***V*** is a matrix of eigenvectors of the respective kernel matrices. We can then express the test statistic in terms of the SVD components as 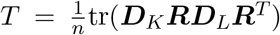, where 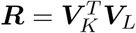 is the outer product between the two eigenvectors. Thus, for a single SNP, the test statistic Is 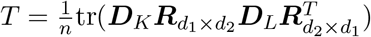, where *d*_1_ = *r* + *s* is the rank of the SQ/CP matrix and *d*_2_ = 1 is the rank of the genotype matrix such that *d*_1_, *d*_2_ ≪ *n*.

#### 4.2.1 Aggregating different LIT implementations using the Cauchy combination test

We explore an important aspect of the test statistic in Equation 5, namely, the role of the eigenvalues in determining statistical significance. The above equations suggest that the eigenvalues of the kernel matrices are emphasizing the eigenvectors that explain the most variation in the test statistic. While this may be reasonable in some settings, the interaction signal can be captured by eigenvectors that explain the least variation and this can be very difficult to ascertain beforehand [40]. In this case, the testing procedure will be underpowered. Thus, we also consider weighting the eigenvectors equally in LIT, i.e., 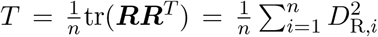, where *D*_R_ are the eigenvalues of the outer product matrix. In this work, we implement a linear kernel (scaled covariance matrix) and so, in this special case, weighting the eigenvectors equally is equivalent to the projection kernel.

In summary, there are two implementations of the LIT framework. The residuals are first transformed to calculate the SQ and CP to reveal any latent interactive effects. We then calculate the weighted and unweighted eigenvectors in the test statistic which we refer to as weighted LIT (wLIT) and unweighted LIT (uLIT), respectively. We also apply a Cauchy combination test (CCT) [41] to combine the *p*-values from the LIT implementations to maximize the number of discoveries and hedge for various (unknown) settings where one implementation may outperform the other. More specifically, let *p*_*c*_ denote the *p*-value for the *c* = 1, 2 implementations. In this case, the CCT statistic is 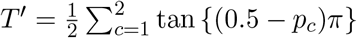, where *π* ≈ 3.14 is a mathematical constant. A corresponding *p*-value is then calculated using the standard Cauchy distribution. Importantly, when applying genome-wide significance levels, the CCT *p*-value provides control of the type I error rate under arbitrary dependence structures. In the Results section, we refer to the CCT *p*-value as aggregate LIT (aLIT).

#### 4.2.2 Incorporating multiple loci in LIT

We can extend LIT to assess latent interactions within a genetic region (e.g., a gene) consisting of multiple SNPs. In the first step, we regress out the joint additive effects from the multiple SNPs along with any other covariates and population structure. In the second step, we calculate the squared residuals and cross products using the corresponding residual matrix. Finally, in the last step, we construct the similarity matrices and perform inference using the HSIC: the linear similarity matrix for the *n×m*_0_ genotype matrix 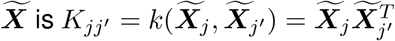 and our test statistic is 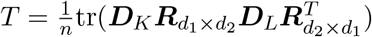 where *d*_2_ = *m*_0_ is the rank of the genotype matrix.

Compared to the previous section, this extended version of LIT is a region-based test for interactive effects instead of a SNP-by-SNP test. A region-based test is advantageous to reduce the number of tests compared to a SNP-by-SNP approach. However, in this work, we demonstrate LIT on SNP-by-SNP genome-wide scan to demonstrate the scalability.

### 4.3 Simulation study

We evaluated the performance of LIT using simulated data with the following assumptions. Let the individual-specific minor allele frequencies of *t* = 1, 2, …, *m* biallelic genotypes be denoted by *π*_*jt*_. Of the *m* SNPs, *m* − 1 SNPs had no interacting partner and a minor allele frequency drawn from a Uniform(0.1, 0.4). The SNP with an interacting partner had a minor allele frequency of 0.25. We fixed this MAF to remove stochastic variation in the observed power induced by simulations differing only by the MAF of the interacting SNP. The genotypes were then drawn from a Binomial distribution with parameter *π*_*jt*_, i.e., *X*_*jt*_ ∼ Binomial(2, *π*_*jt*_). In total, there were *n* = 300,000 individuals simulated to reflect biobank-sized GWAS.

We simulated the trait expression value *Y*_*jk*_ for *k* = 1, 2, …, *r* traits under the polygenic trait model with two risk environmental variables *M*_*j*_ and *W*_*j*_. Specifically, there were *r* = 5, 10 traits and *m* = 100 genotypes simulated with an additive genetic, environmental, and GxE components:

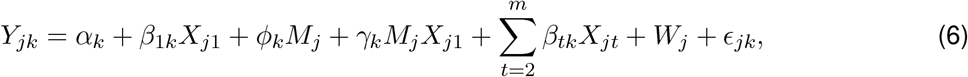

where the intercept, *α*_*k*_, follows a Normal distribution with a standard deviation of 5; the effect sizes of the GxE interaction, *γ*_*k*_, interacting environment, *ϕ*_*k*_, and additive genetic component, *β*_*tk*_, follow a Normal distribution with mean zero and standard deviation of 0.01; the two environmental variables were generated from a standard Normal distribution where only one interacts with the risk allele; and the error term was generated from a standard Normal distribution. Using the above model, we considered different types of pleiotropy. First, we assigned the effect size direction of the additive genetic component, interacting environment, and the GxE interaction to be the same in each trait. We then considered cases where the effect size for the shared GxE interaction is in the same direction (i.e., |*γ*_*k*_|) and random directions across traits. These settings represent positive pleiotropy and a mixture of positive and negative pleiotropy, respectively. We also considered a variation of the above settings where the direction of the effect size for the GxE interaction is opposite of the interacting environment.

We transformed the components in the model using the function 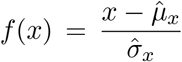, which takes a vector *x* and standardizes it by the estimated mean and standard deviation. We scaled each component to set the baseline correlation between traits (ignoring the risk factor, interactive environment, and GxE interaction) as 0.25, 0.50, and 0.75. In particular, the percent variance explained of the non-interactive environment was 15% and the additive genetic component (minus the risk factor) was 10%, 35%, and 60%, which represents a 0.25, 0.50, and 0.75 baseline correlation between traits, respectively. We then assigned the percent variance explained for the additive genetic risk factor as 0.2%, the interactive environment as a uniformly drawn value from 0.5% to 2.0%, the GxE interaction as a uniformly drawn value from 0.1% to 0.15%, and the remaining variation as noise.

In our simulation study, we also varied the proportion of traits with an interaction term. For *r* traits, let *τ*_*r*_ denote the proportion of traits with a shared GxE interaction signal. We varied this proportion as 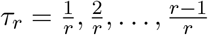. At each combination of baseline trait correlation, number of traits, and proportion of null traits, we generated data from the above polygenic trait model 500 times for each pleiotropy setting. We calculated the empirical power by averaging the total number of times the *p*-values were below a significance threshold of *α* = 5 *×* 10^−8^. Under the null hypothesis of no GxE interaction, we assessed the type I error rate at *α* = 1 *×* 10^−3^ using 50 simulated datasets with 10,000 SNPs where the traits do not have a GxE interaction. We also considered cases where the random error follows a Chi-squared distribution with five degrees of freedom and a *t*-distribution with three degrees of freedom under the null hypothesis.

### 4.4 UK Biobank

The UK Biobank is a collaborative research effort to gather environmental and genetic information from half a million volunteers 40–69 years old in the United Kingdom. The data was collected across 22 assessment centers from 2006 to 2010 where participants were given a general lifestyle and health questionnaire, a physical examination, and a blood test that provided genetic data [64, 65]. See ref. [66, 67] for detailed information on the study design.

We applied LIT to four obesity-related traits, namely, waist circumference, hip circumference, body mass index, and body fat percentage. We restricted our analysis to unrelated individuals with British ancestry and removed any individuals with a sex chromosome aneuploidy. Using the imputed genotypes (autosomes only), SNPs were filtered in PLINK [68] with the following thresholds: a MAF of *>*0.05, a genotype missingness rate of *<*0.05, Hardy-Weinberg equilibrium (defined as *>*10^−5^), and an INFO score of *>*0.9. The traits were adjusted for age and the top 20 principal components provided by the UK Biobank to account for ancestry. We removed individuals with measurements that were four standard deviations above the average and then standardized the traits by sex. After filtering, there were 329,146 individuals and 6,186,503 SNPs in our analysis.

### 4.5 Challenges for latent interaction tests

In the Results section, we addressed a couple of challenges for interaction tests that do not require observing the interactive variable(s). The first is that false positives are possible due to linkage disequilibrium (LD) with a SNP that has a large additive effect (see, e.g., [23]). Intuitively, an imperfect correction of the additive effects creates heteroskedasticity which can be detected after transforming the residuals. Ideally, the SNP(s) in LD with additive effects are regressed out from the traits to avoid false positives. Therefore, we first identified SNPs that were in LD (within 1 Mb and correlation *>*0.1) with the lead SNPs. We then applied a multivariate testing procedure, GAMuT [35], on the selected SNPs to detect additive effects across traits. Note that GAMuT performs an association test between a SNP and the traits using the same test statistic as LIT. The significant SNPs detected were regressed out from the traits and then these adjusted traits were used in LIT.

The second challenge is that an incorrect scaling of the trait, or a scaling where the polygenic assumption does not hold, will induce a variance effect [17, 24]. This complicates underlying inferences because the latent variables may contain non-interactive genetic effects that will be detected as a non-additive effect; for example, the latent variables may capture non-linear effects such as *X*^2^. However, such effects are indistinguishable from a loci with a dominant effect. To be conservative, we addressed the extent of dominance/scaling issues by fitting a two degree of freedom genotypic model to each trait. This allows us to flexibly capture and estimate non-linear genetic variation as a function of genotype. We regressed these effects out from each trait and then applied LIT to the adjusted traits to test whether a SNP remained statistically significant. Under the assumption that a majority of loci act additively, if the significance results across many loci are driven by non-linear genetic effects then it may be suggestive of trait scaling issues.

## Software and data

LIT is publicly available in the R package lit. The package can be downloaded at https://github.com/ajbass/lit. The code to reproduce the results in this work can be found at https://github.com/ajbass/lit_manuscript and access to the UK Biobank data can be requested at https://www.ukbiobank.ac.uk/enable-your-research/apply-for-access.

## Acknowledgements

This research has been conducted using the UK Biobank Resource under Applied Number 58259. This work was supported by NIH grants R01 AG071170 (AJB, SB, DJC, MPE), R01 AG072120 (APW, TSW), R01 AG075827 (APW, TSW), and R01 AG079170 (TSW). The authors would like to thank Michael Boehnke for his helpful feedback during the preparation of this manuscript.

## 5 Appendix

### 5.1 Testing for latent genetic interactions

To review the regression model from the Results section, suppose *Y*_*jk*_ depends on a biallelic locus with genotype *X*_*j*_, an unobserved (or latent) environmental variable *M*_*j*_, and a latent genotype-by-environment (GxE) interaction *X*_*j*_*M*_*j*_ for *j* = 1, 2, …, *n* unrelated individuals with *k* = 1, 2, … *r* measurable traits. The regression model is expressed as

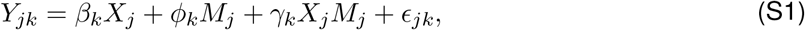

The left side of the equation are the trait values which are observable random variables. The right side contains four components: the observable genotype *X*_*j*_ with effect size *β*_*k*_; an unobservable variable *M*_*j*_ with effect size *ϕ*_*k*_; an unobservable interaction *X*_*j*_*M*_*j*_ with effect size *γ*_*k*_; and an unobservable random error *ϵ*_*jk*_ with mean zero and variance 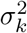. Without loss of generality, we assume that *M*_*j*_ is mean zero with unit variance. Our inference goal is it to test whether *γ*_*k*_ = 0 for *k* = 1, 2, …, *r* without having to observe the latent environmental variable *M*_*j*_.

The following sections are outlined as follows. We first show that a latent genetic interaction induces trait variance and covariance patterns under the above model assumptions. We then review the distributional theory behind the individual-level trait central cross moments. Using these results, we briefly show how latent interactive effects can be detected within a regression model framework.

#### 5.1.1 Latent interactions induce differential variance and covariance patterns

We show in the main text that a latent interaction can be detected based on calculating the individual-specific trait variances (ITV) and covariances (ITC). To construct these quantities, let *e*_*jk*_ = *Y*_*jk*_ − *β*_*k*_*X*_*j*_ denote the trait residuals after removing the additive genetic effect. For simplicity, assume the effect sizes are known. For the *j*th individual, given the genotype *X*_*j*_, the *r × r* individual-specific trait covariance matrix is

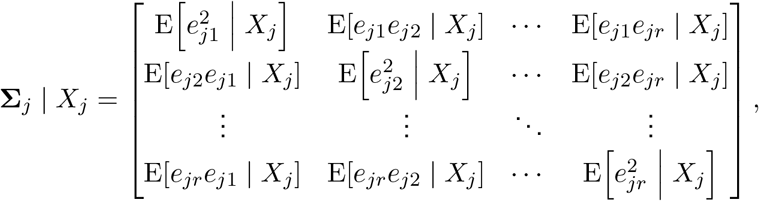

where the ITV are the *r* diagonal elements and ITC are the 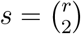 off-diagonal elements.

The presence of a latent interaction shared by multiple traits induces differential ITV and ITC patterns as a function of genotype. More specifically, given our model assumptions, the ITC between the *k*th and *k*^*′*^

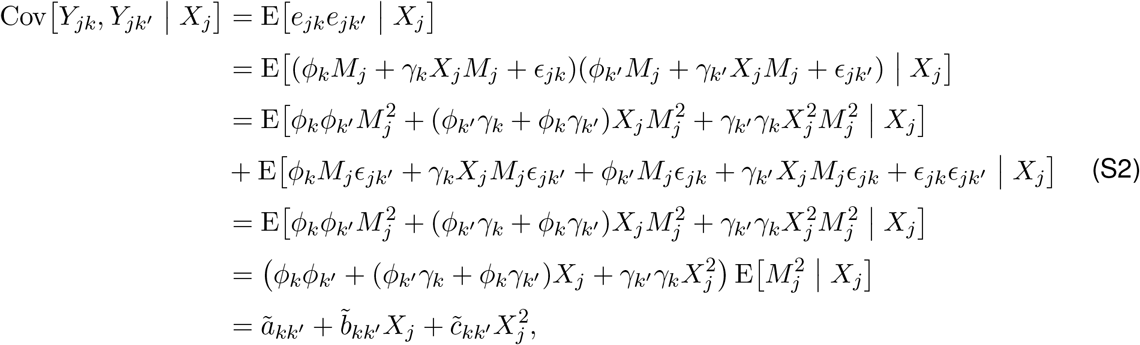

where 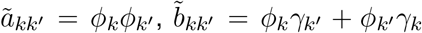, and 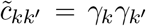. Note that the fourth line follows from our assumption that the random errors of each trait are independent of each other, the genotype, and the environmental variable, and so E[*M*_*j*_*ϵ*_*jk′*_ | *X*_*j*_ = E[*M*_*j*_*ϵ*_*jk*_ | *X*_*j*_] = E [*ϵ*_*jk*_*ϵ*_*jk*_*′*| *X*_*j*_] = 0. The fifth line follows from the assumption that the environmental variable *M*_*j*_ is mean zero with unit variance and independent of the genotype, and so E[*M*_*j*_ | *X*_*j*_] = E[*M*_*j*_] = 0 implying that 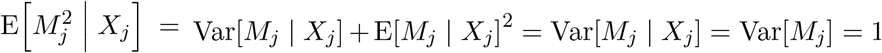. Following similar steps as above, the ITV is

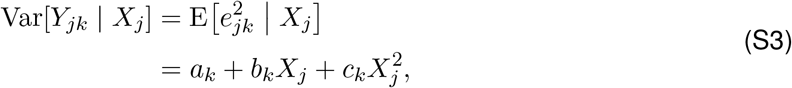

where 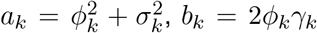 and 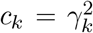. Thus, we have shown that a latent GxE interaction will create differential trait variance and covariance patterns that depend on genotype. In particular, a latent GxE interaction in trait *k* (*γ*_*k*_ *≠* 0) will induce a variance pattern that depends on genotype (Equation S3), and also induce a covariance pattern between traits *k* and *k*^*′*^ when there is a shared interaction (*γ*_*k′*≠_0) or a shared interacting variable (*ϕ*_*k′*_ ≠ 0; Equation S2).

Even though we limit our discussion to a single latent environmental effect and genotype, our results hold more generally under the polygenic trait model. Furthermore, while we consider a simple interaction effect, it is straightforward to show that other complex latent signals involving the genotype induce differential variance and covariance patterns. Although, the exact functional form may be more complicated than above.

#### 5.1.2 Distribution of the cross products

Following the above discussion, we describe the distribution for the cross product of two random variables that follow a Normal distribution. We then use this result to describe the sampling variability of the cross product and squared residual terms within a regression model framework in the next section. To simplify notation, let *Y*_1_ ≡ *Y*_*j*1_ and *Y*_2_ ≡ *Y*_*j*2_ denote the first two traits of the *j*th individual. Without loss of generality, suppose these traits are normally distributed with mean zero, unit variance, and correlation coefficient *ρ*. The cross product term is denoted by *Z* = *Y*_1_*Y*_2_.

The relationship between traits can be expressed as

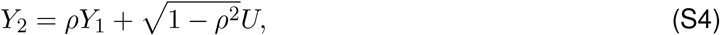

where *U ∼* N(0, 1). The cross product term is then

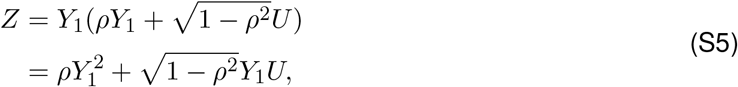

where 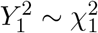 and *Y*_1_*U ∼* B_0_ where B_0_ is the modified Bessel distribution of the second kind of order zero. For perfectly correlated variables, *Z* is distributed as a Chi-squared distribution with one degree of freedom. Alternatively, for uncorrelated variables, *Z* follows a modified Bessel distribution of the second kind of order zero. See ref. [69, 70] for the distribution of the product of two normal random variables.

The first two moments are

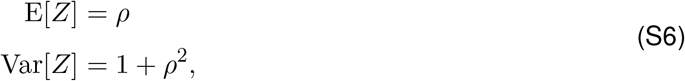

and, more generally, for mean centered traits with variances 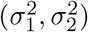, the first two moments are

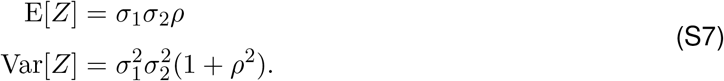

We use this result in the next section to describe the heteroskedasticity in a regression model that treats the cross products or squared residuals as outcome variables.

#### 5.1.3 Regression model for the cross products and squared residuals

Using the central moments result, we first describe the regression model for the cross product terms. Let *P* = *{*(1, 2), (1, 3), …, (2, 3), (2, 4), …, (*r* − 1, *r*)*}* denote the set of cross product pairs such that |*P* | = *s*. The first and second element of the *q*th cross product is *P*_*q*1_ and *P*_*q*2_, respectively, and the cross product between traits is 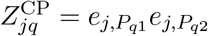. The regression model is

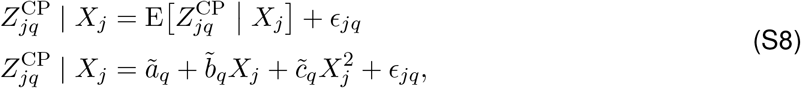

where 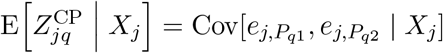 is expressed in Equation S2. The results in Section 5.1.2 can be used to describe the random error in the model: The error term *ϵ*_*jq*_ is independent for *j* = 1, 2, …, *n* observations, but in general, is not normally distributed or identically distributed. Under the null hypothesis of no interactive effects, the errors are identically distributed.

We note that the above regression model differs from typical regression models in two ways. First, the random error does not follow a Normal distribution, although for typical large GWAS sample sizes, this should not impact inference. Second, under the alternative hypothesis where interactions exists, heteroskedasticity arises in the model. To see why, using the results from the previous section, the variance of the error term can be expressed as

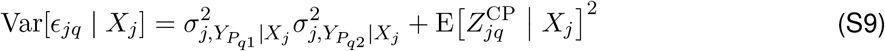

where 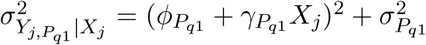 and 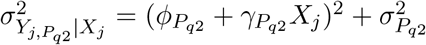. Under the null hypothesis, if the heteroskedasticity is uncorrelated with the explanatory variables then there is type I error rate control. Therefore, controlling for sources of variation such as population structure and nearby SNPs with strong additive effects is important to avoid an inflated type I error rate. Finally, in addition to these sources of variation, an incorrect trait scaling will likely induce heteroskedasticity and also impact type I error rate control.

We briefly state the regression model using the ITV. For the ITV, we are modeling the change in variance of trait *k* as a function of *X*_*j*_:

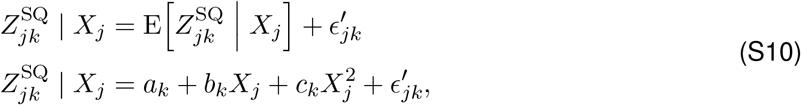

where 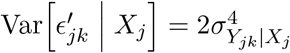. The ITVs are a special case of the ITCs when *ρ* = 1.

Thus far, we assumed that the effect sizes of the additive genetic term is known to simplify the theory. However, in practice, we use the residuals so the above theory does not exactly hold: while the studentized residuals are unbiased estimates, they follow a *t*-distribution and so the squared residuals follow an *F* -distribution (similar adjustments with the cross products). This nuance did not impact any inferences in our simulation study.

There are a few important details with the above regression model approach. First, a test for differential ITV patterns is related to the Breusch-Pagan test [21]. In addition, a regression model on the correlation scale has been discussed elsewhere (see, e.g., [71]) and, more recently, is related to one studied by Lea et al. (2019) [30]. Second, the quadratic relationship between the cross products (or squared residuals) and genotypes only holds for simple interactions, and the underlying (and unknown) functional form is expected to be more complicated. Regardless, for GWAS data where interactions are difficult to detect, *c*_*q*_ (or *c*_*k*_) is likely much smaller than *b*_*q*_ (or *b*_*k*_) and so it is reasonable to assume that the linear term will dominate the signal compared to higher order terms.

### 5.2 Supplementary figures

**Figure S1:**
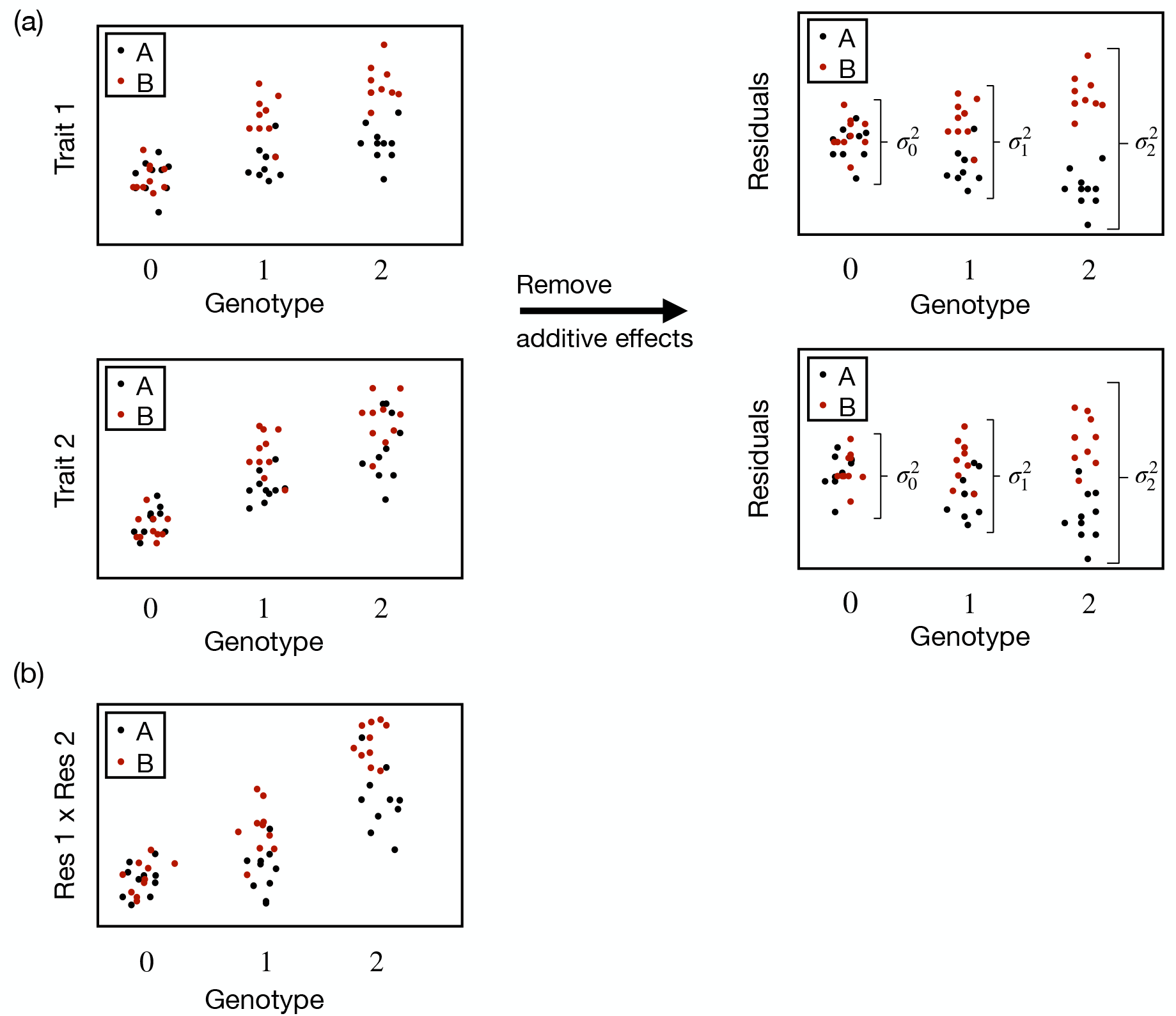
General strategy to detect latent genetic interactions when there are two unobserved environments denoted by ‘A’ and ‘B.’ (a) The additive genetic effect is removed and any heteroskedasticity correlated with genotype implies a latent genetic interaction. (b) When there are two traits measured, the pairwise products between the residuals (cross products) can be used to test for latent genetic effects.

**Figure S2:**
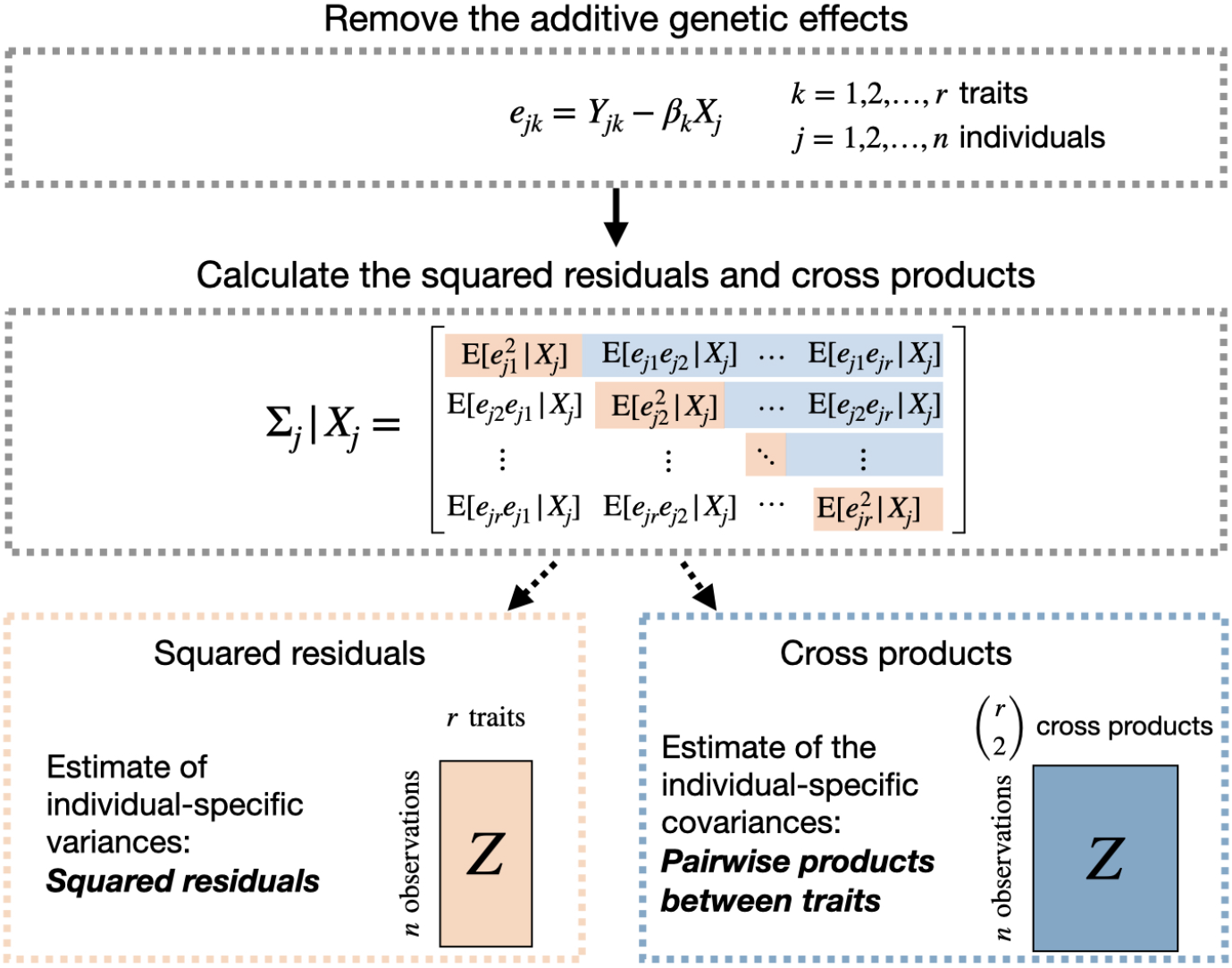
Revealing latent interactive effects using multiple traits. The first step is to remove the additive genetic signal to ensure that the covariance between traits is not caused by the main (additive) effects of the SNP. The individual-specific covariance matrix can then be estimated by calculating the corresponding squared residuals (estimate of the diagonal elements) and the cross products (estimate of the off-diagonal elements). These quantities can be used to infer latent interactive effects.

**Figure S3:**
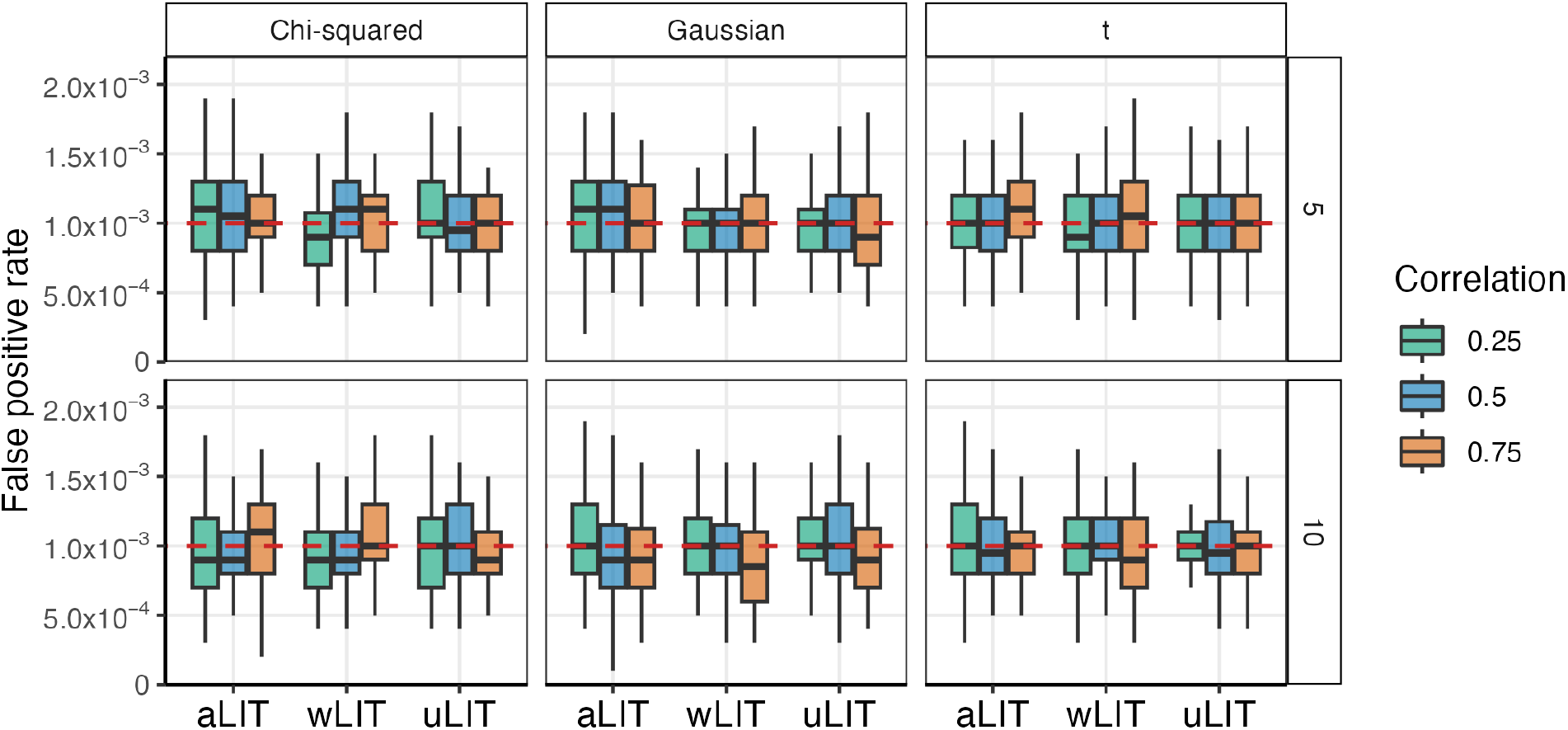
False positive rate of the LIT implementations under the null hypothesis of no interaction. Our simulation study varied the number of traits (rows), baseline trait correlation (0.25 (green), 0.50 (blue), and 0.75 (orange)), and error distribution (columns). For each configuration, there are 50 replicates at a sample size of 300,000. The empirical false positive rate at a type I error rate of 1 *×* 10^−3^ (red dashed line).

**Figure S4:**
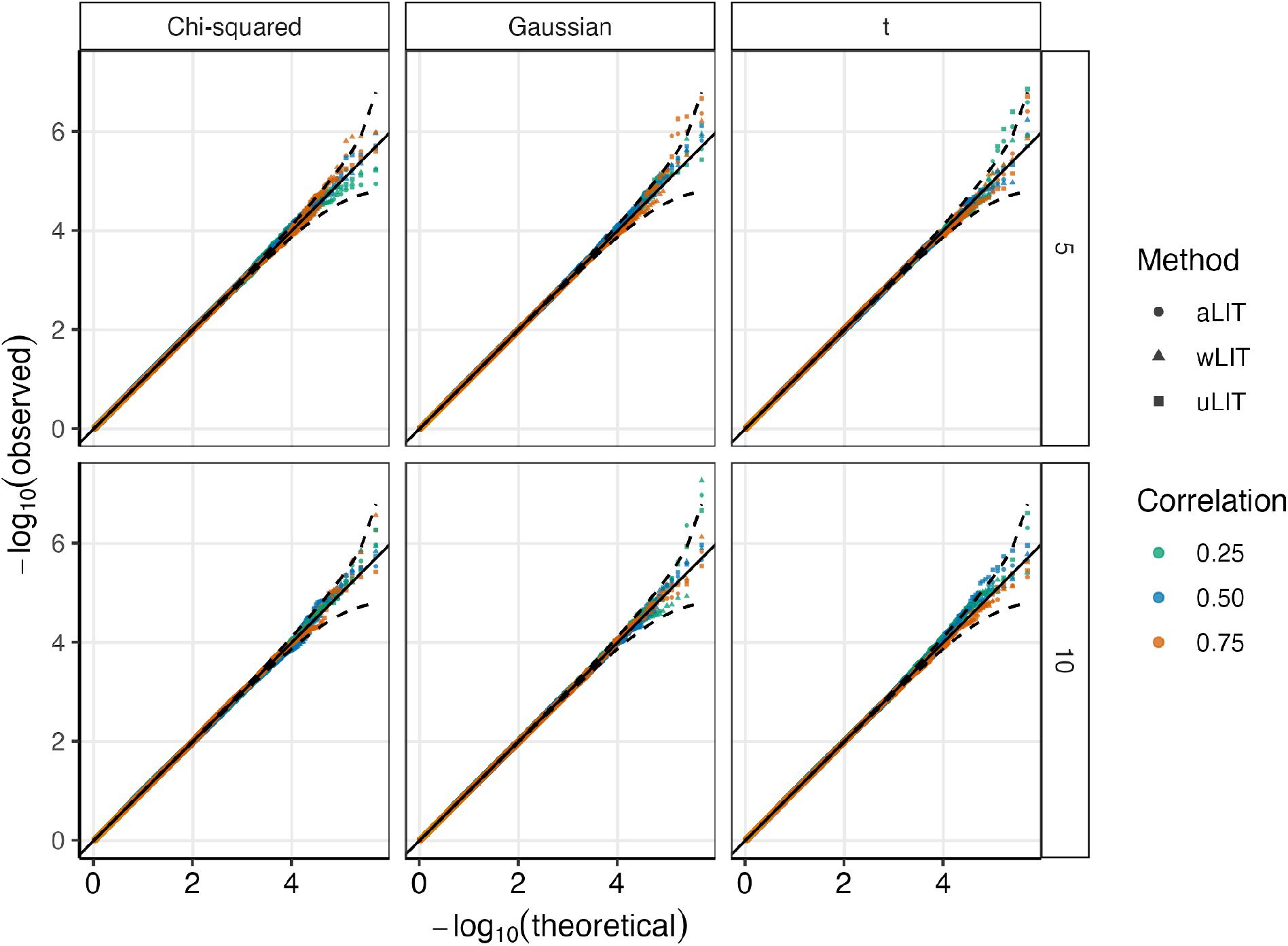
Q-Q plot of the LIT implementations under the null hypothesis of no interaction. Similar to Figure S3, our simulation study varied the number of traits (rows), baseline trait correlation (0.25 (green), 0.50 (blue), and 0.75 (orange)), and error distribution (columns). At each configuration, we simulated 50 datasets of 10,000 SNPs and then combined the *p*-values for a total of 500,000 *p*-values per configuration.

**Figure S5:**
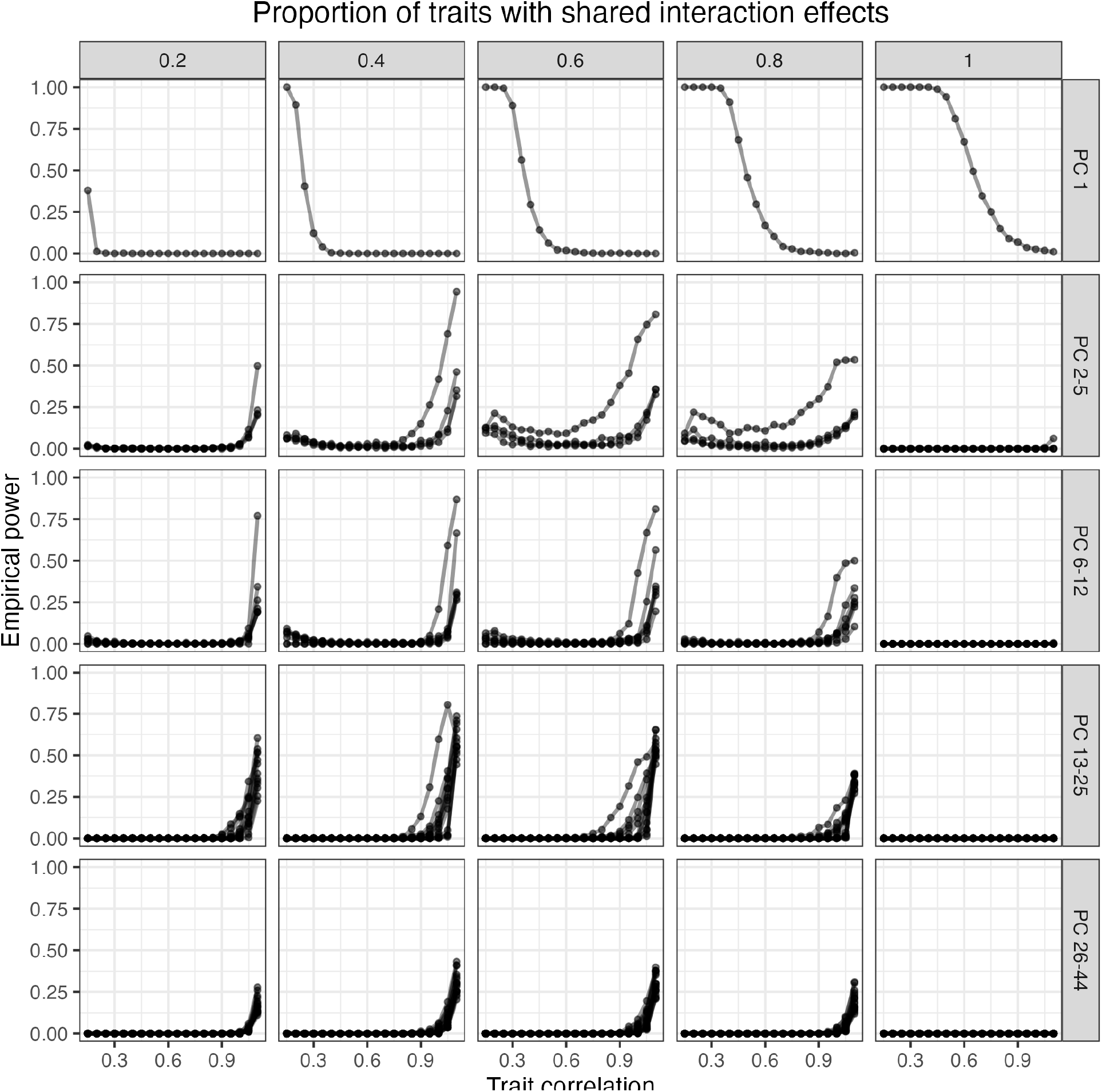
The empirical power of the principal components (rows) for the squared residual and cross product matrix at various baseline correlations (x-axis). In total, there was 10 traits simulated and the proportion of traits with shared interaction effects (columns) was varied. Each point represents the average power across 500 simulations at a significance threshold of 5 *×* 10^−8^.

**Figure S6:**
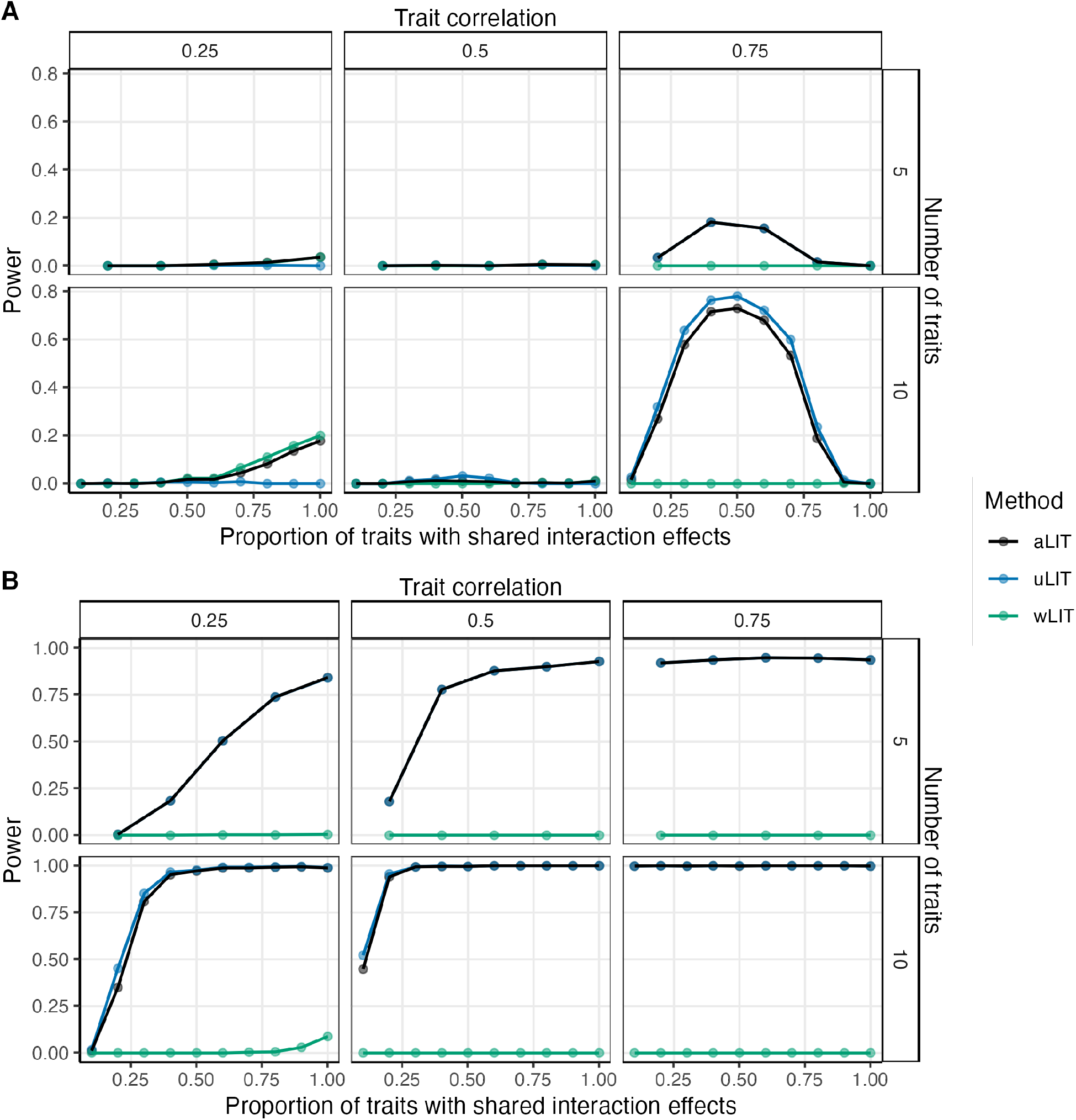
A similar simulation setting to Figure 2 with the direction of the effect size for the interaction term is opposite of the interacting environmental variable under (A) positive pleiotropy and (B) a mixture of positive and negative pleiotropy.

**Figure S7:**
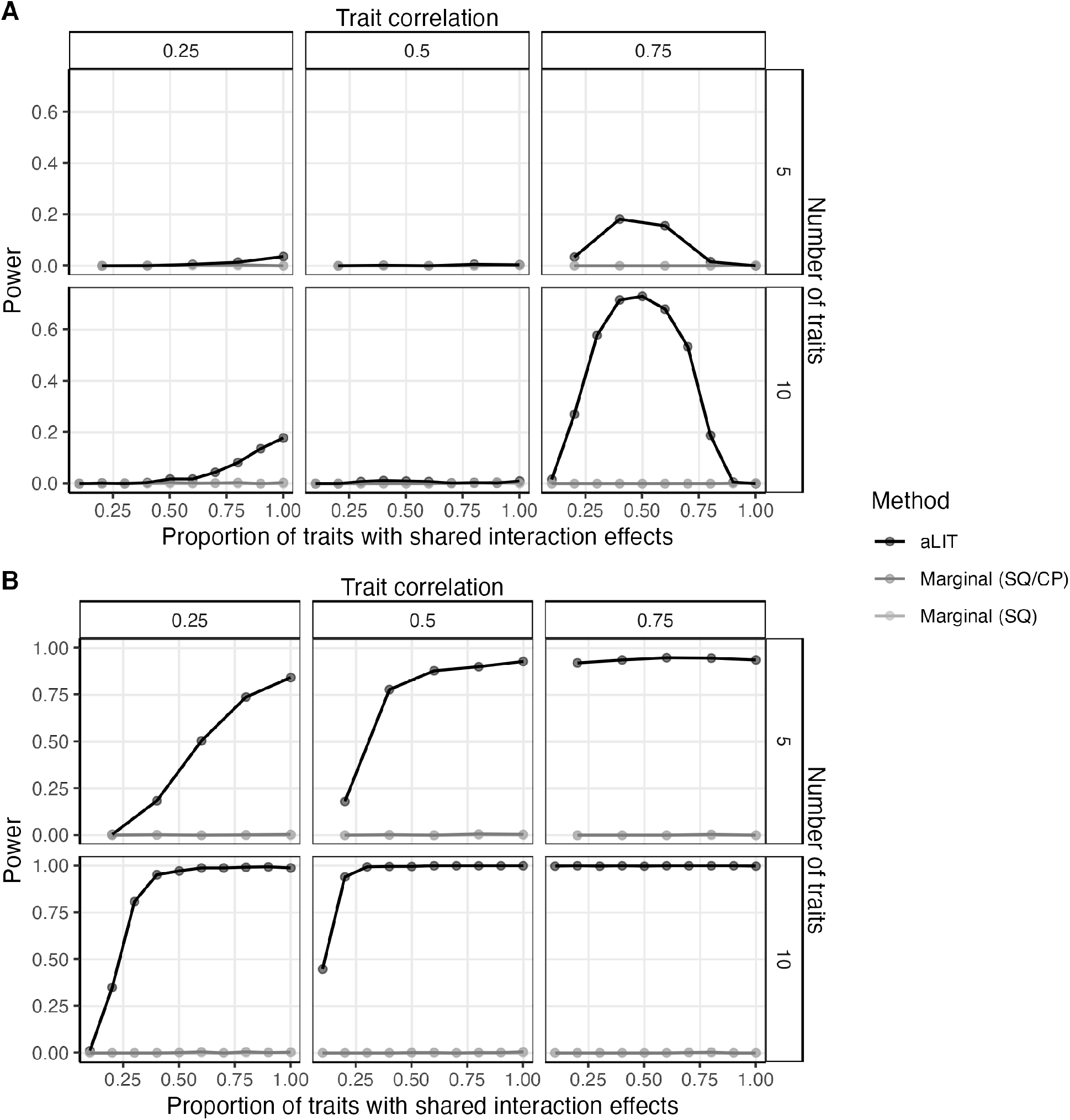
A similar simulation setting to Figure 3 with the direction of the effect size for the interaction term is opposite of the interacting environmental variable under (A) positive pleiotropy and (B) a mixture of positive and negative pleiotropy.

**Figure S8:**
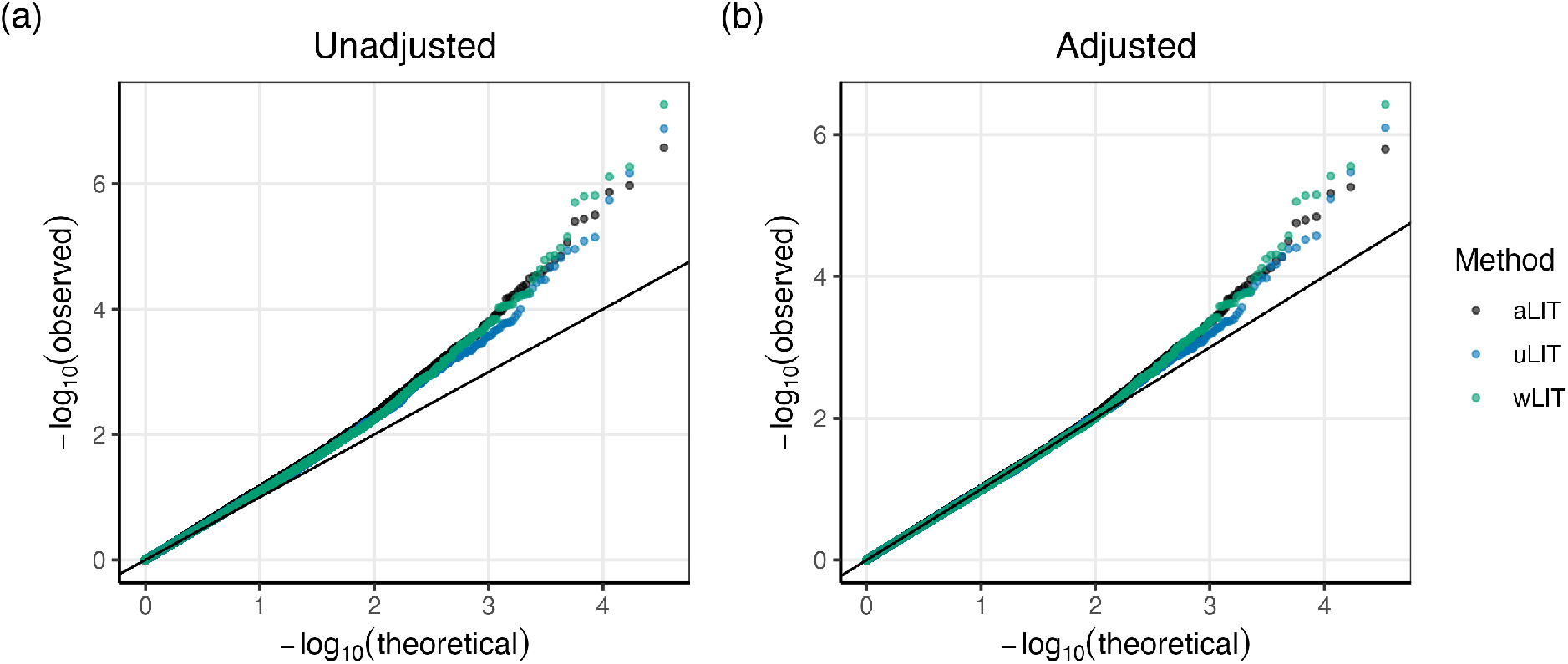
Quantile-Quantile plot of the uLIT, wLIT, and aLIT *p*-values from the UK Biobank. (a) The unadjusted *p*-values and (b) adjusted *p*-values using the genomic inflation factor. The figure removes significant *p*-values and those in strong linkage disequilibrium.

**Figure S9:**
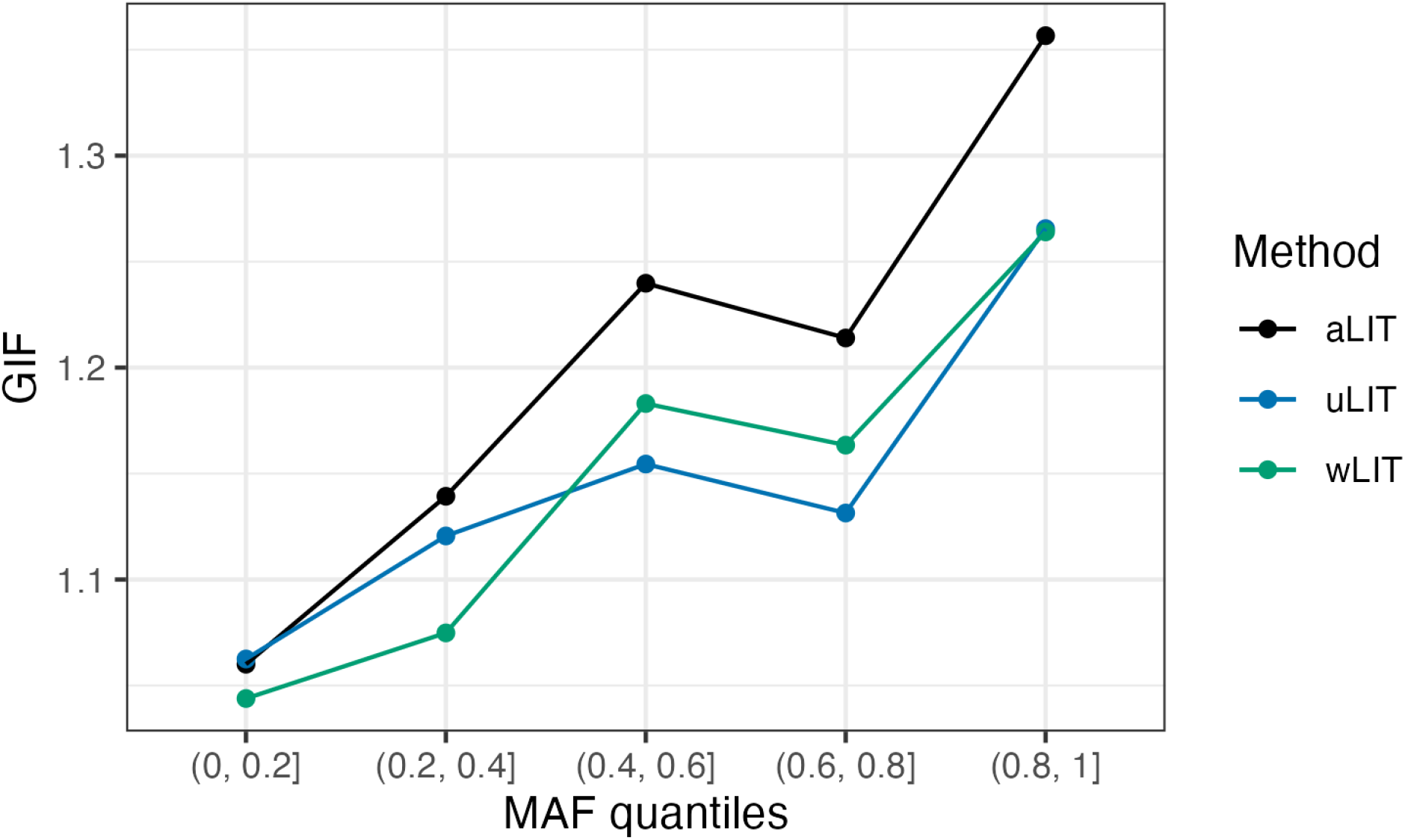
The genomic inflation factor from the UK Biobank analysis using uLIT, wLIT, and aLIT at different minor allele frequency quantiles.

**Figure S10:**
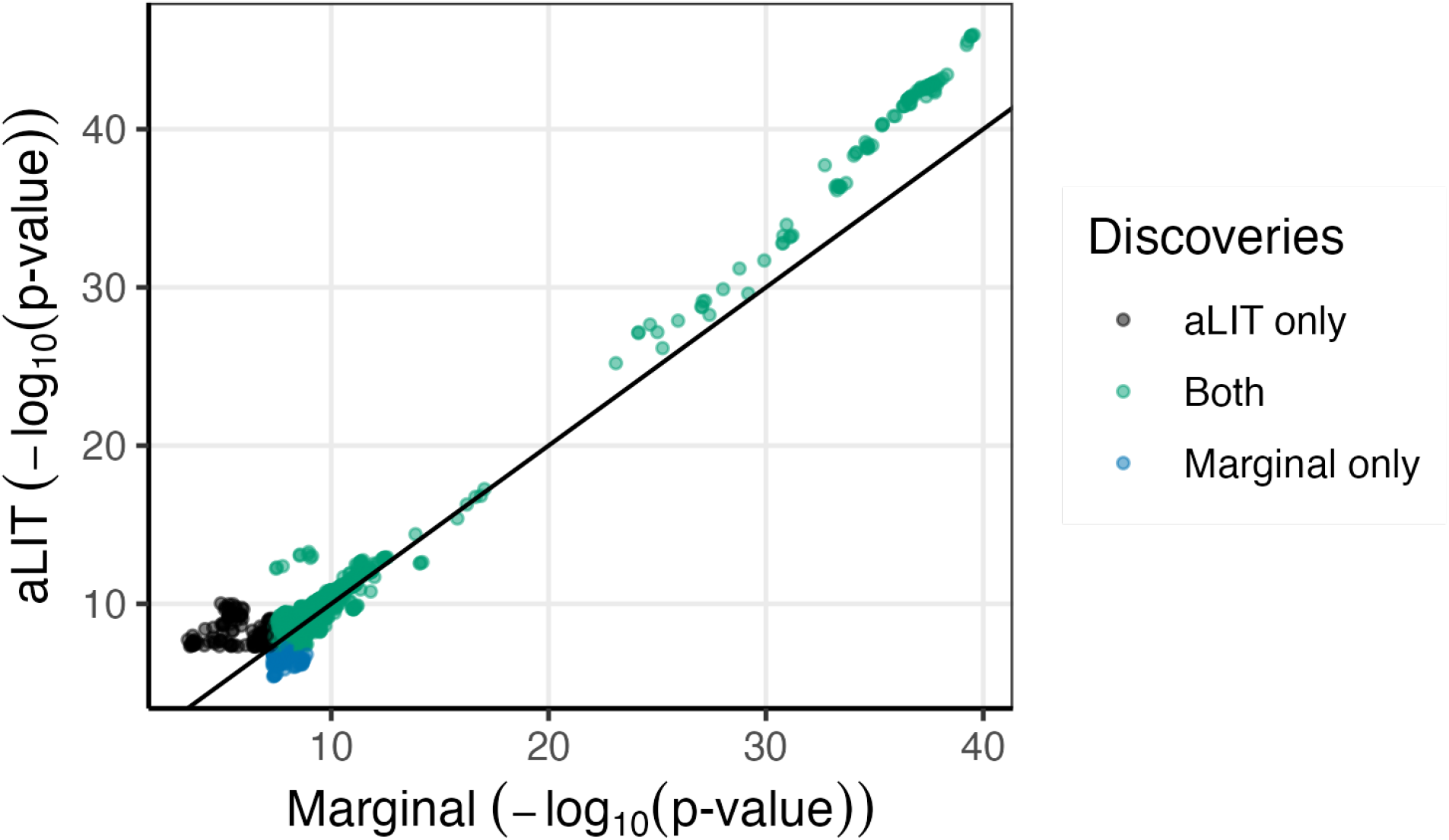
Comparison of the significance results using the marginal testing procedure and aLIT. The genome-wide significance threshold is 5 *×* 10^−8^.

**Figure S11:**
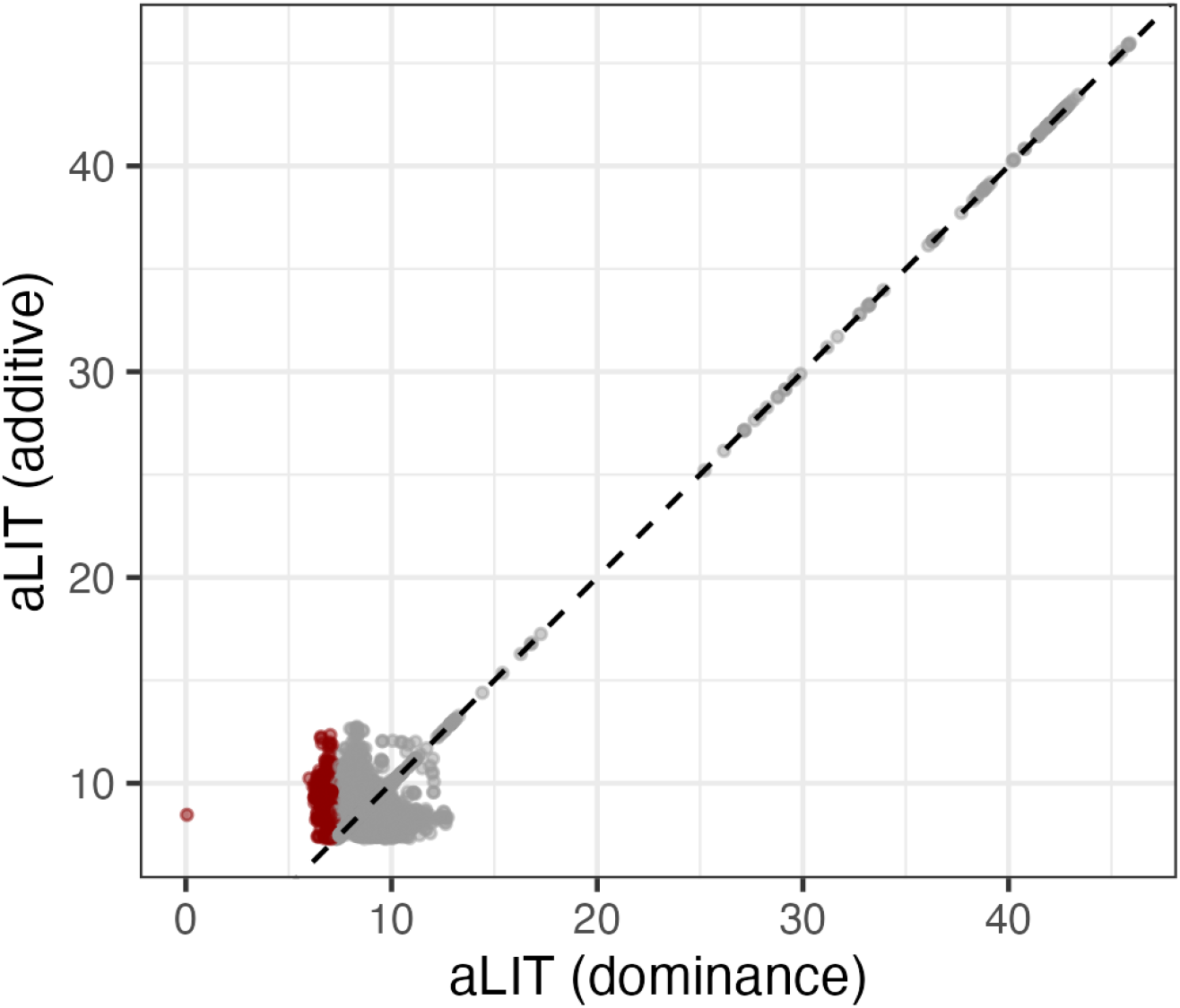
Comparison of aLIT *p*-values after adjusting for additive genetic effects (y-axis) and dominance/scaling effects (x-axis). The dark red points are SNPs that are above the genome-wide significance threshold of 5 *×* 10^−8^. The *p*-values are transformed to be on a logarithmic scale similar to Figure S10.

**Figure S12:**
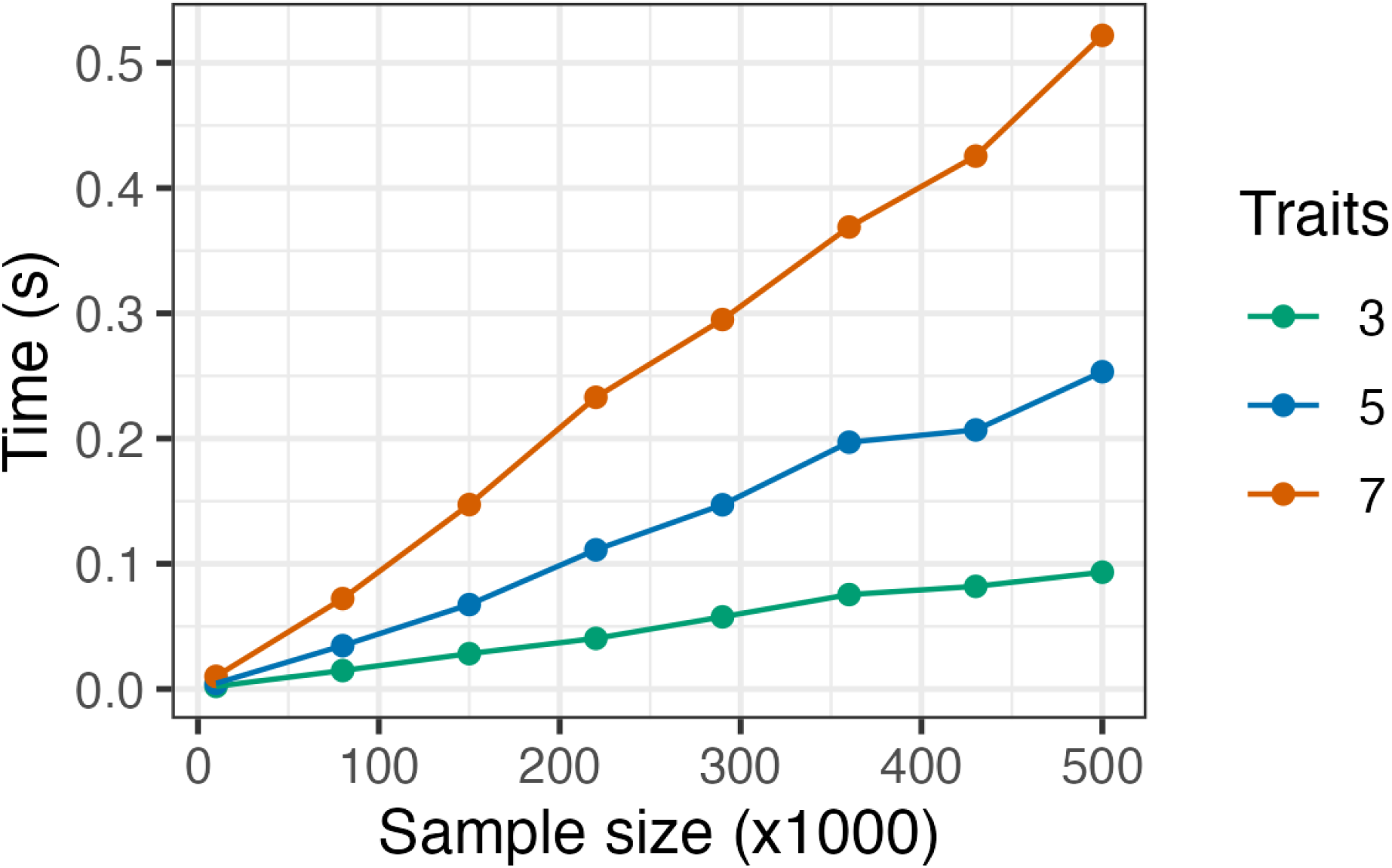
The average computational time to run aLIT on a SNP as a function of sample size and number of traits. Data were simulated the same way in the simulation study and each point is the average time across 500 replicates. Note that only a single core is used and that aLIT can distribute across multiple cores to substantially reduce the computational time.

